# *In Vitro* Platform Establishes Antigen-Specific CD8^+^ T Cell Cytotoxicity to Encapsulated Cells via Indirect Antigen Recognition

**DOI:** 10.1101/2019.12.11.872978

**Authors:** Ying Li, Anthony W. Frei, Ethan Y. Yang, Irayme Labrada-Miravet, Chuqiao Sun, Yanan Rong, Magdalena M. Samojlik, Allison L. Bayer, Cherie L. Stabler

**Affiliations:** J. Crayton Pruitt Family Department of Biomedical Engineering, University of Florida, Gainesville, FL, USA; Graduate Program in Biomedical Sciences, University of Florida, Gainesville, FL, USA; University of Florida Diabetes Institute, Gainesville, FL, USA; Diabetes Research Institute, University of Miami, Miami, FL, USA; Department of Microbiology and Immunology, University of Miami, Miami, FL, USA

**Keywords:** cell replacement therapy, indirect antigen recognition, islet transplantation, immunomodulation

## Abstract

Cell replacement therapy has the potential to cure diseases caused by the absence or malfunction of specialized cells. A substantial impediment to the success of any non-autologous cellular transplant is the need for systemic immunosuppressive drugs to prevent host-mediated rejection of the foreign cells. Cellular encapsulation, i.e., the entrapment of cells within stable polymeric hydrogels, has been clinically explored to prevent host immune recognition and attack, but the efficacy of these encapsulated grafts is poor. While several studies have explored improvements in innate immune acceptance of these encapsulated cells, little attention has been paid to the roles of adaptive immune responses, specifically graft-targeting T cell activation, in graft destabilization. Herein, we established an efficient, single-antigen *in vitro* platform capable of delineating direct and indirect host T cell recognition to microencapsulated cellular grafts and evaluating their consequential impacts. Using alginate as the model hydrogel, encapsulated membrane-bound ovalbumin (mOVA) stimulator cells were incubated with antigen-specific OTI lymphocytes and subsequent OVA-specific CD8^+^ T cell activation and effector function were quantified. We established that alginate microencapsulation abrogates direct T cell activation by interrupting donor-host interaction; however, indirect T cell activation mediated by host antigen presenting cells (APCs) primed with shed donor antigens still occurs. These activated T cells imparted cytotoxicity on the encapsulated cells, likely via diffusion of cytotoxic solutes. Overall, this platform delivers unique mechanistic insight into the impacts of hydrogel encapsulation on host adaptive immune responses, as well as a tool for the efficient immune screening on new encapsulation methods and/or synergistic immunomodulatory agents.

## 1. Introduction

Cell replacement therapy shows great promise in treating diseases caused by the absence or malfunction of specialized cells, such as the replacement of insulin-producing beta cells in Type 1 Diabetes (T1D) or parathyroid cells for parathyroid hormone deficiencies [1–9]. A substantial impediment to the success of any non-autologous cellular transplant, however, is the rejection and/or sensitization of the host to the transplanted cells and the need for systemic immunosuppressive drugs [10, 11]. An alternative to systemic immunomodulation is cellular encapsulation within biocompatible polymers for immunoprotection [12, 13]. The permeability of the encapsulating material is modulated to support the interchange of oxygen, nutrients, and cellular products, while also preventing direct cell interactions between the host and the transplant [14–16]. Numerous materials have been studied for this application, from nanoporous membranes to synthetic hydrogels [13]. Among the numerous choices of materials, alginate, a natural anionic polysaccharide that is easily gelled via exposure to divalent ions at physiological conditions, is one of the most widely used [17]. The encapsulation of various cell types within alginate has resulted in decreased immune rejection and improved long-term graft survival of transplants in rodent models [4, 16–20]. However, the robustness of this protection declines when translated to larger pre-clinical models and clinical trials. For example, the implantation of encapsulated pancreatic islets for the treatment of T1D into non-human primates and humans has been plagued by inconsistent results, with evidence of immune-driven rejection and only modest and transient graft efficacy [21–28].

The failure of alginate encapsulated cells in clinical trials has been attributed to multiple factors, from impurities in the material to inadequate nutrient delivery [12, 29, 30]. Evidence from transplant models also indicates that the host’s adaptive immune cells contribute to loss of graft function, despite retention of the polymeric barrier. For example, xeno-antibodies have been detected following the implantation of alginate encapsulated porcine islets in a nonhuman primate model [23]. Furthermore, others have observed that the addition of T cell co-stimulatory blocking agents (i.e., CTLA4-Ig and anti-CD154 mAb) improves the efficacy of alginate encapsulated cellular grafts in disparate murine and nonhuman primate transplant models [22, 31, 32]. While clear conclusions have not been made to date, due to the multi-factorial nature of these transplant models, these findings indicate that implant-specific adaptive immune responses are activated and may play an active role in the loss of alginate encapsulated grafts.

It has been postulated that the adaptive immune system can detect encapsulated cells via the indirect antigen recognition pathway [12, 33, 34]. For context, immune cells recognize foreign antigens via two pathways: direct and indirect. For direct recognition, host T cells directly recognize antigens presented on the surface of foreign cells via donor MHC molecules [35]. Alternatively, indirect antigen recognition occurs when host antigen presenting cells (APCs) collect and process shed donor antigens for presentation to host T cells in a host MHC-restricted manner [35–37]. Regardless of the antigen presentation pathway, the final consequence is the clonal expansion and activation of graft-specific effector immune cells (*e.g.*, cytotoxic CD8^+^ T cells, helper CD4^+^ T cells, and B cells) that destroy the foreign grafts. For polymeric encapsulation, it has long been hypothesized that direct antigen recognition is prevented by the polymer barrier. However, indirect antigen recognition may still occur due to the shedding of foreign antigens through the biomaterial into the peri-transplant site, as summarized in Figure 1.

**Figure 1.**
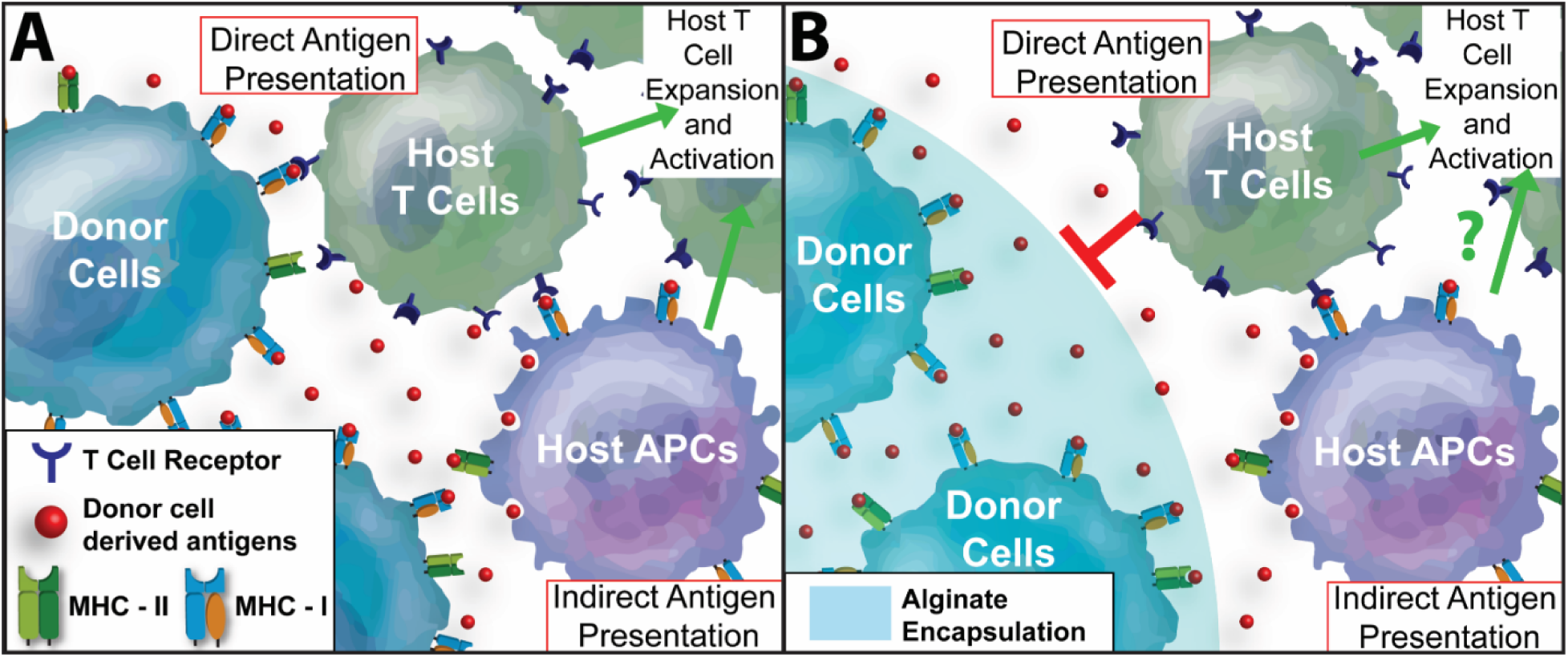
Summary of Proposed Pathways for Antigen Recognition of Unencapsulated and Microencapsulated Cells by the Host Adaptive Immune System. (A) For unencapsulated cell grafts, both direct and indirect antigen recognition pathways occur. For direct antigen recognition, host CD4^+^ and/or CD8^+^ T cells detect the offending foreign antigens presented directly on the transplanted tissue or cells, via donor MHC-II and MHC-I molecules, respectively. Indirect antigen recognition to unencapsulated cell grafts can also occur via host antigen-presenting cells (APCs) presenting donor antigen to host CD4^+^ and/or CD8^+^ T cells via host MHC molecules. Following antigen presentation, T cells proliferate and activate effector functions. (B) For encapsulated cells, e.g. alginate microencapsulation, direct donor antigen presentation to host T cells should be blocked by the polymer barrier. However, antigens shed by the donor cells can diffuse out of the hydrogel for collection and presentation by host APCs, leading to indirect antigen recognition and subsequent antigen-specific T cell expansion and activation.

Despite this long-standing hypothesis, the immune crosstalk between host T cells and the encapsulated graft remains unclear. Without a more comprehensive understanding of the pathways of host T cell activation and its subsequent role, if any, in damaging the underlying encapsulated cells, the capacity to generate immunoprotective biomaterial approaches for long-term clinical efficacy will remain elusive. While animal transplant models can provide physiologically relevant insight into adaptive immune responses to encapsulated grafts, the multitude of other factors that contribute to graft failure, such as the transplant environment, insufficient nutrient delivery, and foreign body responses, overlay and prevent clear delineation of their respective roles. Furthermore, reliance on animal models to characterize these responses is expensive and time-consuming.

Herein, we developed a reproducible, efficient, and antigen-specific *in vitro* platform to conduct mechanistic studies into the role of polymeric microencapsulation on host adaptive immune responses. To impart antigen specificity, encapsulated cells were sourced from membrane-bound ovalbumin (mOVA) mice. These stimulator cells isolated from mOVA transgenic mice express ovalbumin on their cell membrane and present OVA peptide derivatives as self-antigen in MHC-I [38]. To examine OVA-specific adaptive immune activation, responder cells were isolated from OT1 mice, a transgenic murine strain with a dominant CD8^+^ T cell population expressing an engineered alpha/beta T cell receptor recognizing OVA_257-264_ peptide (*i.e.*, SIINFEKL). Combining alginate encapsulation and these antigen-specific immunoreactive cells in a co-culture platform permitted the exploration and delineation of the roles of direct versus indirect antigen recognition and their impact on host T cell activation. This platform was then validated using microencapsulated mOVA pancreatic islets, where the corresponding OT1 T cell activation and its subsequent immune impact on the encapsulated cells was investigated. The contribution of adaptive immune cell recognition and activation on the destabilization of encapsulated transplants is also discussed.

## 2. Methods and Materials

### 2.1 Animals

All animal procedures were conducted under IACUC approved protocols at the University of Florida. Responder cells were sourced from OT1/GFP mice, 8-15 weeks of age, where all OTI CD8^+^ T cells specifically recognize SIINFEKL via transgenic T cell receptors. Stimulator cells (splenocytes and islets) were sourced from mOVA mice (C57BL/6J (CAG-OVA)916Jen/J; Jackson laboratories), whereby all beta-actin expressing cells express both membrane-bound OVA and OVA peptide derivatives as self-antigen in MHC-I [38].

### 2.2 Splenocyte isolation and islet isolation

Spleens were collected from donors in cold HBSS buffer (Corning). A single-cell suspension was prepared by mechanically rupturing the spleen and filtering through a 40-µm cell strainer (Corning). Splenocytes were harvested by spinning down at 500×g for 5 min and erythrocytes were removed by 5 min treatment with ACK lysing buffer. Islets were isolated from mOVA or C57BL/6J mice as previously described [39, 40]. Isolated islets were cultured at 37°C, 5% CO_2_ in a humidified incubator in CMRL 1066 media (Mediatech) supplemented with 10% FBS (Hyclone, GE Healthcare), 20□mM HEPES buffer, 100 U/mL penicillin□streptomycin, and 2□mM L□glutamine until used for experiments.

### 2.3 Encapsulation materials and cell encapsulation

Sterile 1.6% (w/v%) UP MVG (cGMP grade, Pronova, NovaMatrix) alginate in saline was used as the encapsulating material. Splenocytes were encapsulated using a parallel air flow bead generator (Biorep, Inc.) with an average air rate of 3300 mL/min and 50 mM BaCl-MOPS crosslinking buffer, as previously reported [41]. Encapsulation density was 5×10^7^ or 1×10^7^ cells/mL alginate, depending on the experimental design. After three PBS washes, splenocyte microbeads were maintained in complete T cell media (RPMI 1640 medium supplemented with 10% FBS, 100 U/mL penicillin□streptomycin, 2□mM L□glutamine, 20 mM HEPES buffer and 55 μM β-mercaptoethanol). For islet encapsulation, islets were counted 24 hrs post-isolation using a standard algorithm for the calculation of 150□µm diameter islet equivalent (IEQ) number [42] and encapsulated at a density of 1000 IEQ/mL alginate. Encapsulated islets were maintained in CMRL media overnight prior to experimentation. Cell-free alginate microbeads were generated as a control. Bead size was measured (n ≥ 10 for each encapsulation) with Carl Zeiss™ Primo Vert™ Inverted Microscope.

### 2.4 OTI/GFP splenic cell purification by cell sorting

Freshly isolated OTI/GFP splenocytes were labelled with anti-mCD3-Pacific Blue; anti-mCD8a-APC; anti-mCD4-PE/Cy5 and anti-mCD11c-PE antibodies at 4 °C for 20 min (**Table 1**). Labeled cells at the concentration of 25×10^6^/mL were sorted using BD FACSAria™ sorter with over 98% efficiency. i) GFP+CD3+CD11c-CD8a+CD4-cells were sorted out as viable OTI splenic CD8^+^ T cells; and ii) GFP+CD3-CD11c+CD8a+CD4-cells were harvested as the viable OTI splenic cross-presenting CD8a^+^ DCs. Sorting was conducted by the Interdisciplinary Center for Biotechnology Research Cytometry Core and the Center for Immunology and Transplantation at the UF. A representative sorting scheme is shown in **Figure S1**.

### 2.5 mOVA-OTI *in vitro* co-culture platform

To delineate direct and indirect antigen recognition pathways, 1) 100,000 purified OTI CD8^+^ T cells; 2) 75,000 sorted OTI CD8^+^ T cells with 25,000 pre-primed cross-presenting CD8a^+^ DCs (with or without inhibition); or 3) 100,000 unsorted OT1 splenocytes were used as immune responders and co-incubated with 100,000 mitomycin C (50 µg/mL) pretreated unencapsulated or encapsulated mOVA stimulator cells for 48 hrs. All immune responders were labeled with CellTrace™ violet dye for proliferation tracking. The frequency of proliferating granzyme B expressing OTI CD8^+^ T effector cells was quantified as the readout of the co-culture system. Unstimulated control (T cell media), anti-CD3/CD28 Dynabeads^®^ (ThermoFisher); 0.1μM SIINFEKL (InvivoGen) peptide, 0.1μM soluble ovalbumin (InvivoGen) protein, and cell-free alginate microbeads were used as control stimulators for OTI T cell activation. The stimulator/responder (S/R) ratio of the co-culture system was adjusted, as outlined in the experimental design. For mOVA islet-OTI responder co-culture, 50 mOVA or C57BL/6J islets, in unencapsulated or alginate encapsulated form, were hand-picked and co-cultured with 100,000 CellTrace™ violet-labeled OTI/GFP splenocytes at 37°C for 72 hrs before downstream analysis. The same control groups were applied as mentioned above. For cross-presentation inhibition, 25,000 fresh sorted viable CD8a^+^ DCs were primed with 100,000 unencapsulated or encapsulated mOVA cells for 6 hrs with or without 5μg/mL brefeldin A (Sigma)[43]. 75,000 Cell Trace™ Violet labeled purified OTI CD8^+^ T cells were then added for 48 hrs stimulation.

### 2.6 Flow cytometry

To evaluate the level of OVA-specific CD8^+^ T cell activation, OTI responders were collected after co-culture for flow cytometry analysis. OTI responders were sequentially stained with Live/Dead^®^ Fixable Near IR dye (Invitrogen), anti-mCD8a-PE, anti-mCD25-PE/Cy7, and anti-mGranzymeB-APC (**Table 1**) for viability assessment and immune phenotyping. Background signals were identified and excluded by isotype-matched and fluorescence-minus-one controls. The level of OTI CD8+ T cell activation was quantified as the percentage of proliferating granzyme B+ CD8^+^ T effector cells (**Figure S2**). Data were acquired using BD LSRII or FACSCelesta analyzer with a proper compensation setting. Data analysis was performed using FCS Express 6.05 software (DeNovo software). Proliferation index (PI, the total number of cell divisions) and division index (DI, the average number of cell divisions) were calculated by fold dilution of CellTrace™ Violet signal and proliferation modeling using the embedded module of FCS Express 6.05 software.

### 2.7 *In vitro* islet assessments

To evaluate mOVA islet viability and function after co-culture experiment, Live/Dead^®^ imaging and static glucose-stimulated-insulin-release (GSIR) assay were used. For Live/Dead^®^ assay, the encapsulated islets post co-culture were stained with 26.67 μM calcein AM and ethidium homodimer-1 at 37°C for 30 min. Confocal images were then acquired using Leica TCS SP8 microscope. Fluorescent signals from calcein AM and ethidium homodimer-1 were quantified as the area of calcein AM+ (or ethidium homodimer-1+) normalized by the area of an islet using ImageJ software (n ≥ 10 images per group per test). For the GSIR assay, samples with 50 encapsulated islets were immobilized in chromatography columns using Sephadex G10 resin beads (GE Health) after co-culture, as previously described [40]. The columns were then sequentially stimulated with 1 hr of low (3 mM) glucose (L1), followed by 1 hr of high (16.7 mM) glucose (H), and lastly an additional 1 hr of low (3 mM) glucose (L2). Samples (1 mL) collected after each 1-hr stimulation were analyzed for insulin content via ELISA (Mercodia) and normalized by the islet number (50) of each sample.

### 2.8 Statistical analysis

Data are expressed as the mean ± standard deviation for each group, with individual data points shown as scattered points. For all experiments, a minimum of three independent biological replicates were included in each group. Outlier screening was performed for all data sets using Robust Fit with Cauchy estimate with multiplier K=2, using SAS JMP Pro v13.1.0. software. Statistical assessments were performed using one-way ANOVA with Tukey’s multiple comparison analysis using GraphPad Prism Software. Statistical differences were considered significant when the probability value (*p*) was <0.05. Statistical difference is shown as — *\*p<0.05*; *\*\*p<0.01*; *\*\*\*p<0.001*; *\*\*\*\* p < 0.0001* and *n.s.* indicates *not significant*.

## 3. Results

### 3.1 Alginate encapsulation blocks contact-dependent direct antigen recognition

To provide an efficient means to evaluate host adaptive immune responses to encapsulated cells, an OVA-based antigen-specific platform was designed. Stimulator cells sourced from mOVA spleens were encapsulated within alginate microbeads formulated from ultra-pure, cGMP-grade, medium viscosity alginate (UP-MVG). Responder lymphocytes were sourced from OTI mice, which contain a high percentage of CD8^+^ T cells clonally specific to OVA derived SIINFEKL peptide [44]. To characterize CD8^+^ T cell activation, the frequency of viable, proliferating, and granzyme B^+^ CD8^+^ cells was quantified via flow cytometry, as these markers classically define an effector cytolytic T cells (**Figure S2**) [45, 46]. For select experiments, proliferation analysis of CD8^+^ T cells was also conducted.

To examine the role of direct antigen recognition in microencapsulation systems, CD8^+^ T cells were sorted from OTI splenocytes (**Figure S1**). Resulting sorted OTI CD8^+^ T cells were incubated with either unencapsulated or encapsulated mOVA cells and the activation of OTI CD8^+^ T cells was subsequently measured (Figure 2A). The response of purified OTI CD8^+^ T cells to unencapsulated mOVA cells was efficient and robust, with high activation (90.5 ± 1.3%) and proliferation (PI = 3.12 ± 0.23) (Figure 2B&D); the activation profile was statistically equivalent to CD8^+^ T cell responses to anti-CD3/28 activator beads (*p=0.05*, Tukey post-hoc), demonstrating vigorous stimulation. Contrarily, OT1 CD8^+^ T cell proliferation and activation was completely ablated when co-cultured with alginate encapsulated mOVA cells; T cell activation (2.7 ± 2.2%) and proliferation (PI = 1.35 ± 0.31) was statistically comparable to unstimulated controls (*p=0.96*, Tukey post-hoc; Figure 2C&D). While *in vitro* cultures of primary splenic T cells typically become apoptotic in the absence of stimulation [47], T cells harvested from co-cultures of both unencapsulated and encapsulated mOVA cells were highly viable (*p<0.0001* vs unstimulated control respectively; Tukey post-hoc) albeit statistically distinct (*p = 0.005*; t-test; Figure 2D inset & **S3A**).

**Figure 2.**
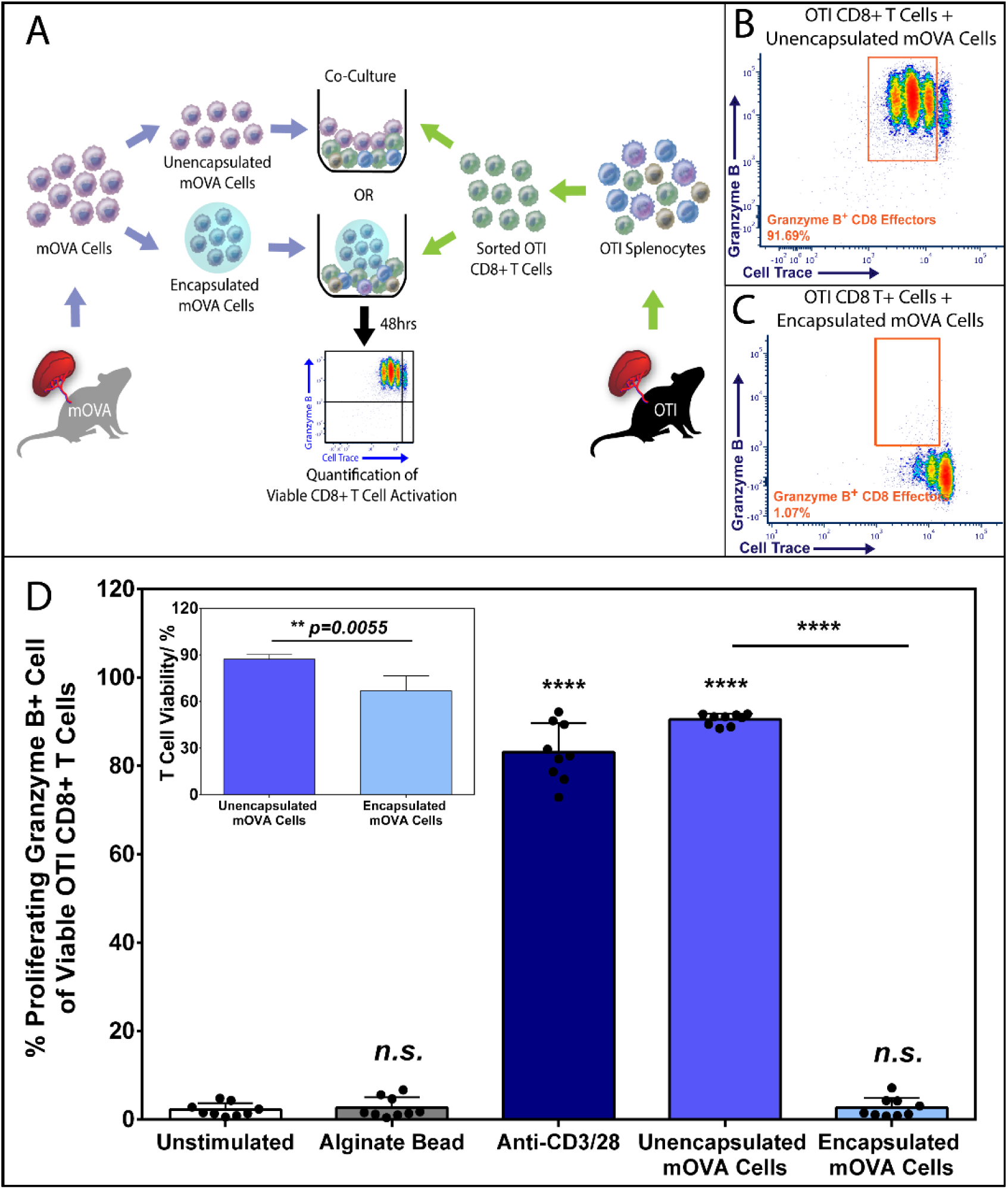
Alginate Encapsulation Blocks Cell-Contact Dependent Direct CD8^+^ T Cell Antigen Recognition and Activation. (A) Schematic overview of the 48hr co-culture experiment co-incubating sorted OTI CD8^+^ T cells (see **Figure S2** for sorting) with unencapsulated or encapsulated mOVA stimulatory splenocytes at 1:1 ratio. Antigen-specific OTI CD8^+^ T cell activation was quantified by the % of proliferating (Cell Trace^®^ Violet labeled) and granzyme B + CD8+ T effector cells (see **Figure S1** for gating) via flow cytometry analysis. Representative FCM data of OTI CD8^+^ T cell activation by unencapsulated (B) or alginate encapsulated (C) mOVA cells after a 48-hour stimulation. (D) Summary of the frequency of proliferating granzyme B+ CD8^+^ T effectors in response to designated stimuli (x-axis). Inset: CD8^+^ T cell viability after 48 hrs incubation with unencapsulated or encapsulated mOVA cells. Bars indicate the average of individual data points (N=3; n=9) with standard deviation. Statistical significance was determined as *\*\*\*\*p < 0.0001 and n.s. = not significant* via Tukey’s test when compared with the unstimulated control group.

To validate the inert nature of the base alginate biomaterial, cell-free alginate microbeads were included as a control group. CD8^+^ T cell responses to this group were statistically equivalent to unstimulated controls (*p=0.87*; Tukey post-hoc; Figure 2D). Of note, endotoxin levels from alginate eluates were below assay detection limits (sensitivity 0.1EU/mL, **Table 2**).

To confirm the antigen specificity of the co-culture system, cells sourced from C57BL/6J mice were screened as non-specific stimulators (**Figure S4A**). OTI CD8^+^ T cells were nonresponsive to either unencapsulated or encapsulated C57BL/6J splenic cells; the percentage of effector CD8^+^ T cells were statistically equivalent to unstimulated controls (*p = 0.54 and 0.97*, respectively; Tukey post-hoc). As OTI mice are syngeneic with the C57BL/6J strain [38], the lack of T cell response within the same time frame of this assay validates that the previously observed OTI CD8^+^ T cell activation in response to mOVA stimulators was OVA antigen-specific.

### 3.2 Indirect antigen recognition is preserved in alginate microencapsulation

Following the conclusion that alginate encapsulation suppresses direct antigen recognition, we sought to develop methods to assess activation via indirect antigen recognition pathways. While the CD8^+^ T cell is the dominant responsive population of lymphocytes in OTI splenocytes due to the nature of its transgenic development [48], the splenocytes of OTI mice also contains CD4^+^ T cells, CD11c^+^ dendritic cells (DC), F4/80^+^ macrophages, and B cells **(Figure S5)**. These immune cells can play a role in activating cytolytic CD8^+^ T cells via indirect antigen presentation pathways, such as through DC cross-presentation and CD4^+^ T cell licensing [49, 50]. Thus, it was postulated that the inclusion of the full OTI splenic repertoire in this co-culture platform could permit the study of CD8^+^ T cell activation via indirect antigen mechanisms (Figure 3A).

**Figure 3.**
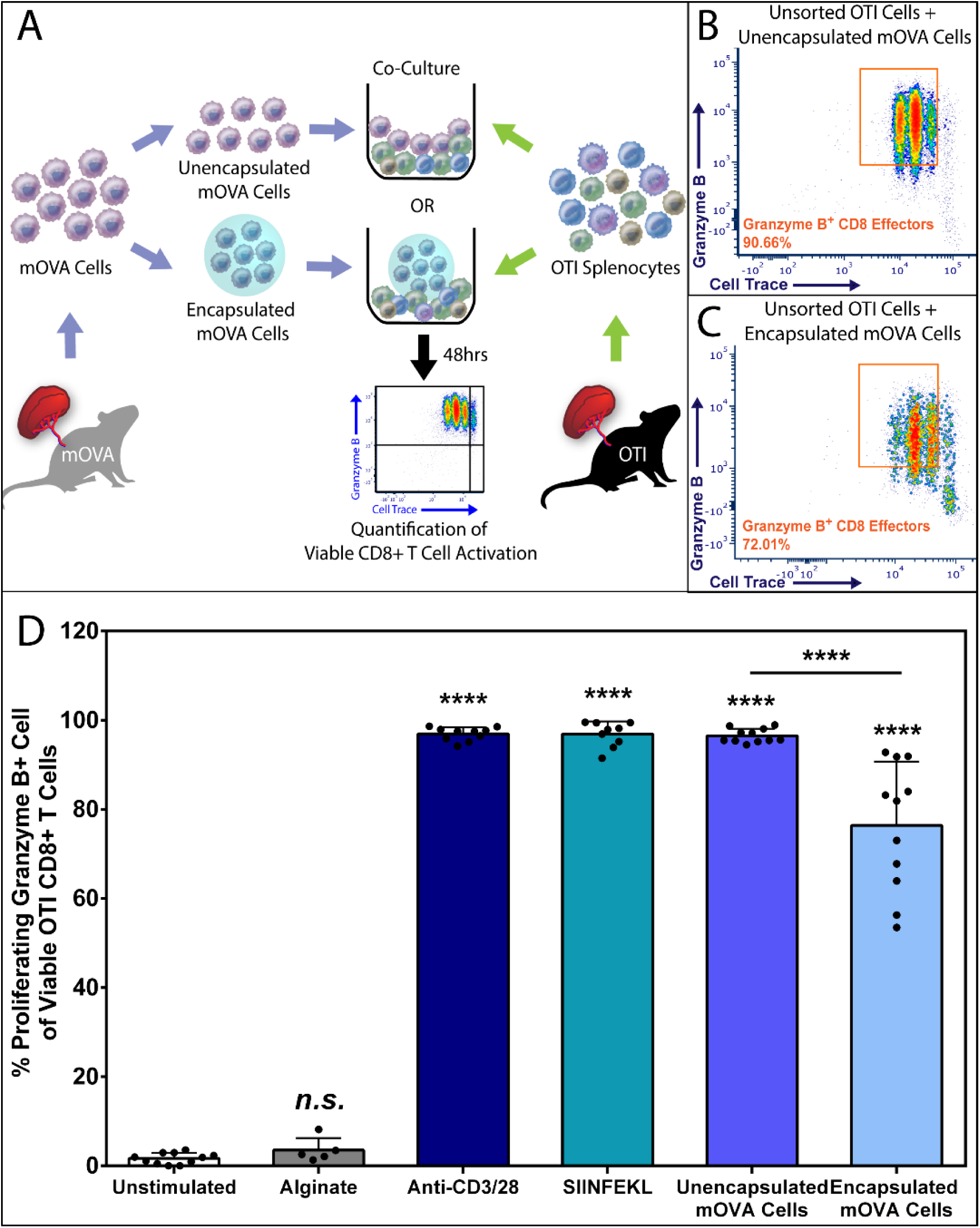
Alginate Encapsulated Cells Effectively Activate Antigen-Specific CD8^+^ T cells via the Indirect Antigen Recognition Pathway. (A) Schematics of the 48hr co-culture experiment co-incubating mOVA stimulator cells, either unencapsulated or encapsulated form, with the unsorted OTI splenic immune responder cells. OVA-specific OTI CD8^+^ T cell activation was quantified by the % of proliferating and granzyme B + CD8^+^ T effectors via flow cytometry analysis (see **Figure S1**). Representative FCM data of OVA-specific CD8^+^ T cell activation within the unsorted OT1 responder pool in response to equivalent amount of unencapsulated (B) or alginate encapsulated (C) mOVA cells. (D) Summary of the frequency of proliferating granzyme B+ CD8^+^ effector T cells, gated from the whole OTI splenocyte population, in response to designated stimuli (x-axis). Bars indicate the average with individual data points (N=4; n=6-11) shown with standard deviation. Statistical difference was determined as *\*\*\*\*p < 0.0001; n.s. = not significant* via Tukey’s test when compared with the unstimulated control group.

As expected, the co-culture of OTI splenocytes with unencapsulated mOVA cells resulted in significant CD8^+^ T cell activation (96.5 ± 1.5%) and proliferation (PI = 3.63 ± 0.52); values were statistically equivalent to cells stimulated via polyclonal activator beads (*p>0.999*, Tukey post-hoc) or soluble SIINFEKL peptide (*p>0.999*, Tukey post-hoc) (Figure 3B&D). Distinct from purified CD8 T cell cultures, when whole OTI splenocytes were incubated with encapsulated mOVA cells, the resulting CD8^+^ T cell activation and expansion was surprisingly vigorous, with significant activation (76.4 ± 14.3%; *p < 0.0001* vs unstimulated control) and proliferation (PI = 2.30 ± 0.60), albeit statistically lower than those measured in response to unencapsulated mOVA cells (*p<0.0001*; Figure 3C&D). This robust CD8^+^ T cell activation to alginate encapsulated mOVA cells, particularly when compared to the complete inhibition of CD8+ activation for purified CD8+ T cell responders, indicates that the inclusion of endogenous splenic lymphocytes alters T cell activation pathways. Of note, similar to the results from purified CD8^+^ T cell studies, stimulators incubated with either cell-free alginate microbead controls (Figure 3D) or C57BL/6J cells (**Figure S4B**) exhibited T cell activation equivalent to unstimulated controls.

### 3.3 Professional APCs are mediators of indirect antigen recognition to alginate encapsulated cells

It is known that CD8a^+^ dendritic cells (DC) uptake, process, and cross-present extracellular antigens to CD8^+^ T cells via the exogenous MHC-I pathway, resulting in the generation of antigen-specific CD8^+^ effector T cells [50]. Based on the distinct OTI CD8^+^ T cell activation profile in response to the alginate encapsulated mOVA cells when comparing whole splenic responders to the purified CD8^+^ T cells, it was hypothesized that host APCs facilitate CD8^+^ T cell in the indirect activation given a polymeric barrier prevents direct T cell contact. To investigate this theory, OTI CD8a^+^ cross-presenting DCs were sorted from bulk OTI splenic responders via flow cytometry, as outlined in **Figure S1**. Sorted DCs were then incubated with mOVA stimulating cells for a six-hour antigen priming, followed by the addition of responding OTI CD8^+^ T cells (Figure 4A). To further delineate the role of cross-presentation for indirect T cell activation, brefeldin A, an inhibitor of cross-presentation [43], was added to select groups during antigen priming.

**Figure 4.**
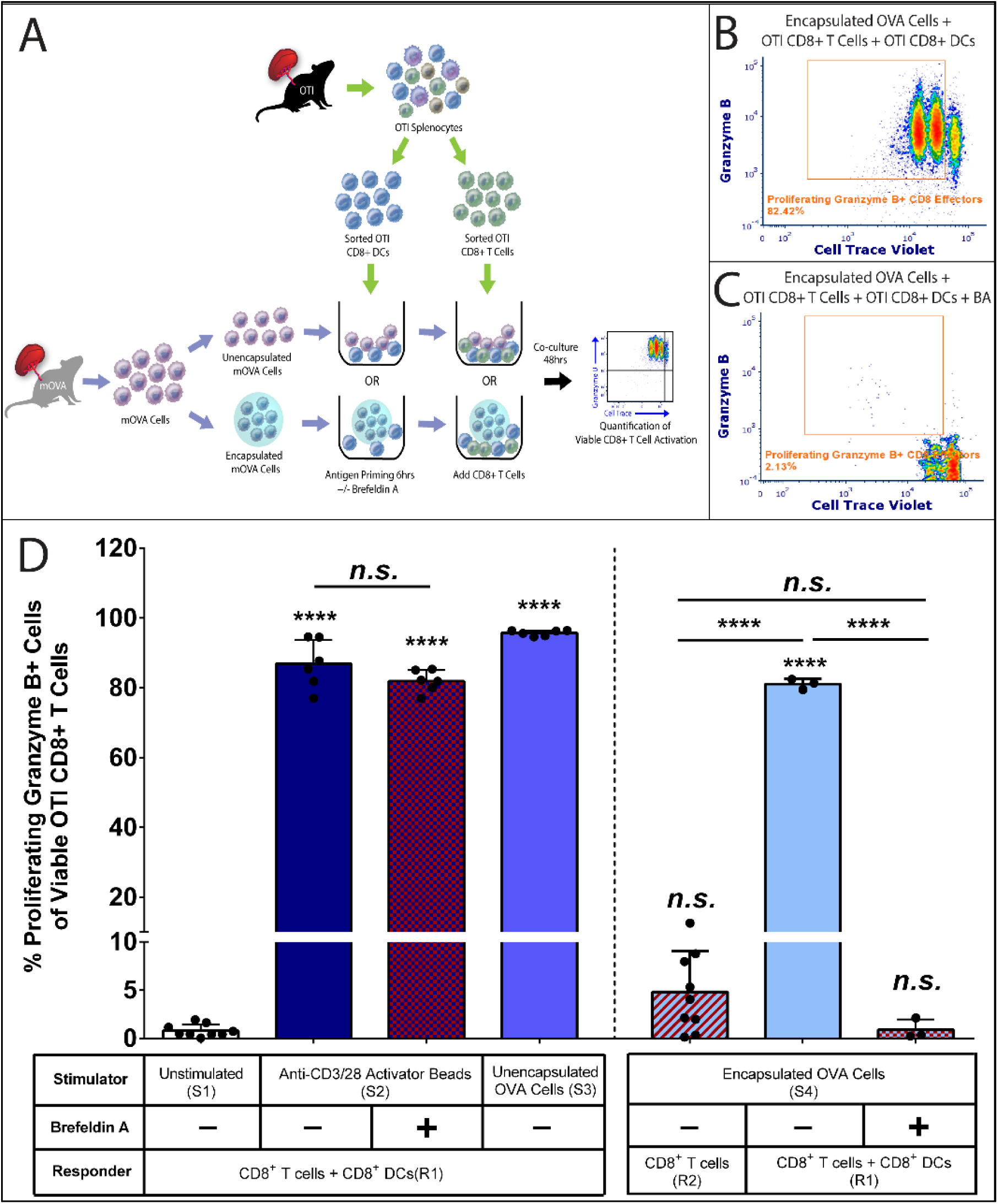
Cross Presenting CD8+ Dendritic Cells Contribute to Indirect CD8^+^ T cell Activation Instigated by Alginate Encapsulated Cells. (A) Schematics of the co-culture experiment. Firstly, 15000 purified cross-presenting CD8^+^ DCs (see **Figure S2**) were primed by either unencapsulated or encapsulated mOVA cells for 6hrs with or without the cross-presentation inhibitor brefeldin A (BA, 5μg/mL). Then 85,000 purified CD8^+^ T cells (see **Figure S2**) with Cell Trace^®^ Violet labeling were added to the system for a 48hr co-culture. OVA-specific OTI CD8^+^ T cell activation was quantified by the % of proliferating and granzyme B + CD8^+^ T effectors via flow cytometry analysis (see **Figure S1**). Representative FCM data of OTI CD8^+^ T cell activation within the CD8^+^ T cell and CD8^+^ DC responder pool stimulated by encapsulated mOVA cells (B) with or (C) without brefeldin A treatment. (D) Summary of the frequency of proliferating granzyme B+ CD8^+^ effector T cells in response to designated stimuli (table at the bottom). Stimulator groups are annotated as unstimulated control (S1); anti-CD3/28 activator beads (S2); unencapsulated mOVA cells (S3) and encapsulated mOVA cells (S4). Responder groups are annotated as CD8^+^ T cells + CD8a^+^ DCs (R1) and purified CD8^+^ T cells (R2). Cross-presentation is labeled as untreated (BA-) or inhibited (BA+). Bars indicate the average with individual data points (n=3-9) shown with standard deviation. Statistical difference was determined as *\*\*\*\*p < 0.0001; n.s. = not significant* via Tukey’s test when compared with the unstimulated control group or in between groups.

As summarized in Figure 4D, the inclusion of purified CD8a^+^ DCs into CD8^+^ T cell cultures generated targeted effects. Consistent with our previous findings, robust CD8^+^ T cell activation was observed within CD8^+^ T cells and CD8a^+^ DCs (responder group R1) were co-cultured with activator beads (anti-CD3/28, group S2 + R1) (86.8 ± 7.0%) or unencapsulated mOVA cells (group S3 + R1) (95.7 ± 0.8%). When encapsulated mOVA cells were used as stimulators (S4), CD8^+^ T cell response was altered depending on the composition of the OT1 responder pool. Specifically, if purified CD8^+^ T cells were applied as immune responders, minimal activation was observed (4.8 ± 4.2%; group S4 + R2), consistent with the results reported in Section 3.1. However, the inclusion of CD8a^+^ DCs into the co-culture system substantially altered this response, resulting in robust CD8^+^ T cell activation (81.0 ± 1.2%; group S4 + R1; Figure 4B&D) statistically equivalent to that observed in response to anti-CD3/28 activator beads (*p = 0.27*; Tukey post-hoc). To delineate the role of cross-presenting CD8a^+^ DCs in the indirect CD8^+^ T cell activation, brefeldin A (BA) inhibition was applied to the co-culture system. The inclusion of brefeldin A during DC antigen priming resulted in the potent loss of the downstream indirect OT1 CD8^+^ T cell activation (1.2 ± 0.9%; group S4 + R1 + BA; Figure 4C&D), with a granzyme B negative, nonresponsive CD8^+^ T cells population (Figure 4C); the resulting activation level was comparable to purified CD8^+^ T cell controls (*p = 0.74*; group S4 + R2, no BA; Tukey post-hoc). Of note, brefeldin A inhibition for the control anti-CD3/28 activator bead stimulation (group S2 + R1 + BA) did not impair CD8^+^ T cell activation (*p = 0.08*; Tukey post-hoc) or viability (**Figure S3C**, *p=0.56*; Tukey post-hoc), validating that brefeldin A treatment does not compromise the activation capacity of CD8^+^ T cells in this presented model.

## 3.4 Indirect T cell activation level correlates to encapsulated cell number and density

To further characterize the dynamics of T cell activation in response to encapsulated cells, a two-parameter antigen titration was performed, whereby the overall encapsulated stimulator cell number, as well as encapsulation density, were independently varied, followed by the quantification of the subsequent T cell activation accordingly. Whole OTI splenic responders and experimental parameters used for the titration were identical to those defined in Figure 3A.

As summarized in Figure 5A, antigen dose dependency was observed for CD8^+^ T cells stimulated by either unencapsulated or alginate encapsulated mOVA cells. The trend of the increased OTI CD8^+^ T cell activation in response to unencapsulated mOVA cells was less evident, with robust stimulation observed using a modest stimulating cell dose (*i.e.*, 10,000 mOVA). Contrarily, antigen dosage from encapsulated mOVA stimulators strongly influenced the response of CD8^+^ T cells, with activation values ranging from insignificant to high (*e.g.*, 2.8 ± 0.1% and 48.5 ± 0.8% in response to 10,000 and 50,000 mOVA encapsulated cells, respectively). Comparing CD8^+^ T cell responses to titrated levels of unencapsulated or encapsulated mOVA cells, it can be concluded that the presence of the alginate barrier plays a suppressive role in T cell activation. For example, the degree of CD8^+^ T cell activation in response to 50,000 encapsulated mOVA cells was almost half (51.8%) of its unencapsulated counterpart. To further dissect the difference in direct and indirect CD8^+^ T cell responses, proliferation modeling was employed (Figure 5B). A decrease in proliferating CD8^+^ T cell number (5260 ± 781) and a delay in T cell proliferation (PI = 1.78 ± 0.04) was measured for CD8^+^ T cells responding to encapsulated mOVA cells, when compared its matched unencapsulated counterpart (T cell number: 10614 ± 1321, *p< 0.0001* and PI: 3.22 ± 0.05, *p < 0.0001*, respectively; Tukey post-hoc). However, the immunosuppressive effect is incomplete, with increased indirect CD8^+^ T cell activation measured when the antigen dose increased (Figure 5A).

**Figure 5.**
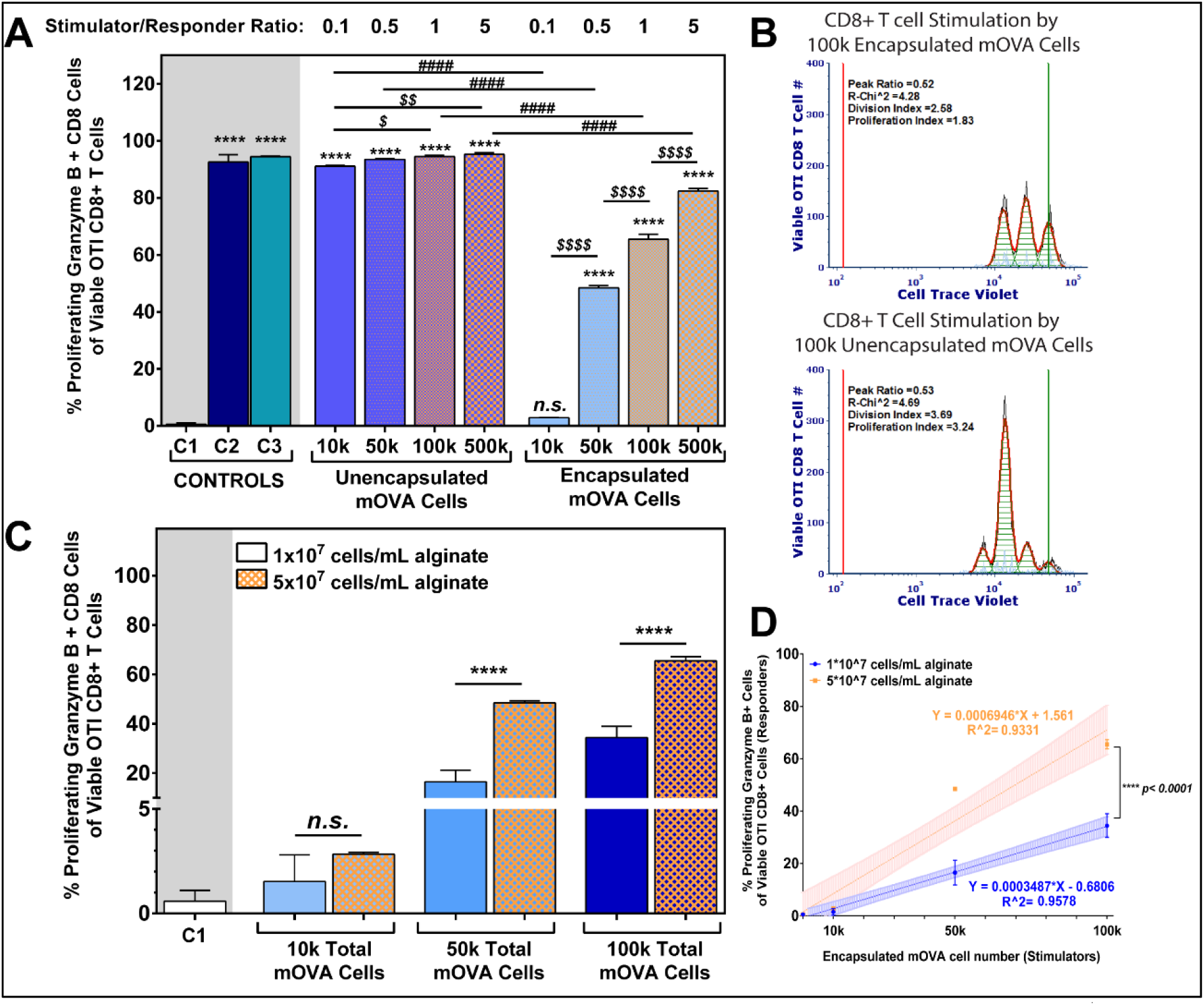
Distict Antigen Dosage Impacts for Unencapsulated versus Alginate Encapsulated. (A) OTI CD8^+^ T cell activation in response to titration of unencapsulated (red bars) and encapsulated (blue bars) mOVA stimulator cells in a 48hr co-culture (n=3); S/R ratio ranged from 0.1 to 5 (top label). Loading density for encapsulated cells at 5×10^7^ cells/mL alginate. Controls: C1: unstimulated; C2: anti-CD3/28 activator beads; C3: 0.1μM SIINFKEL peptide. Mean comparisons to unstimulated control C1 (statistics shown as *); within unencapsulated or encapsulated mOVA stimulators (statistics shown as $); and between unencapsulated and encapsulated stimulators (statistics shown as #) using Tukey’s test. (B) Representative proliferation modeling of OTI CD8^+^ T cells activation in response to 100,000 unencapsulated or encapsulated mOVA cells. (C) OTI CD8^+^ T cell activation in response to titration of alginate encapsulated mOVA cells with two different cell densities (1×10^7^ or 5×10^7^ cells/mL alginate; top legend; n=3). Means were compared using Tukey’s test. Error = standard deviation. (D) Linear correlation between OTI CD8^+^ T cells activation levels and encapsulated mOVA cell number with two different densities (1×10^7^ in blue or 5×10^7^ in orange cells/mL alginate; top legend), with the regression equations shown and coefficient of determination labeled as R^2. Shaded area = standard deviation. Slope comparison analysis was performed between the regression curves. Statistical difference was determined as *\* p< 0.05; ** p< 0.01; ****p < 0.0001; n.s. = not significant*.

Of interest, the cell density within the microencapsulated grafts also played a role in modulating the level of CD8^+^ T cell activation. Specifically, increasing the cell density within the encapsulation platform consistently resulted in amplified responder cell activation, despite an equivalent total cell dosage (Figure 5C). Figure 5D summarizes this phenomenon, where the two cell loading densities resulted in distinct activation trends (*p<0.0001*, slope comparison, t-test).

### 3.5 Alginate encapsulated pancreatic islets activate T cells via indirect antigen presentation pathway

To examine the host adaptive immune responses to encapsulated cells that are relevant to a disease model, pancreatic islets were screened. In this approach, pancreatic islets were isolated from mOVA mice and encapsulated within alginate microbeads, followed by co-culture with whole OTI splenic responders. Antigen-specific CD8^+^ T cell activation in response to mOVA islets was evaluated (Figure 6A). For stringent capture of the indirect T cell activation, the complete encapsulation of the antigenic islet was visually validated and islet microbeads were handpicked for all experiments.

**Figure 6.**
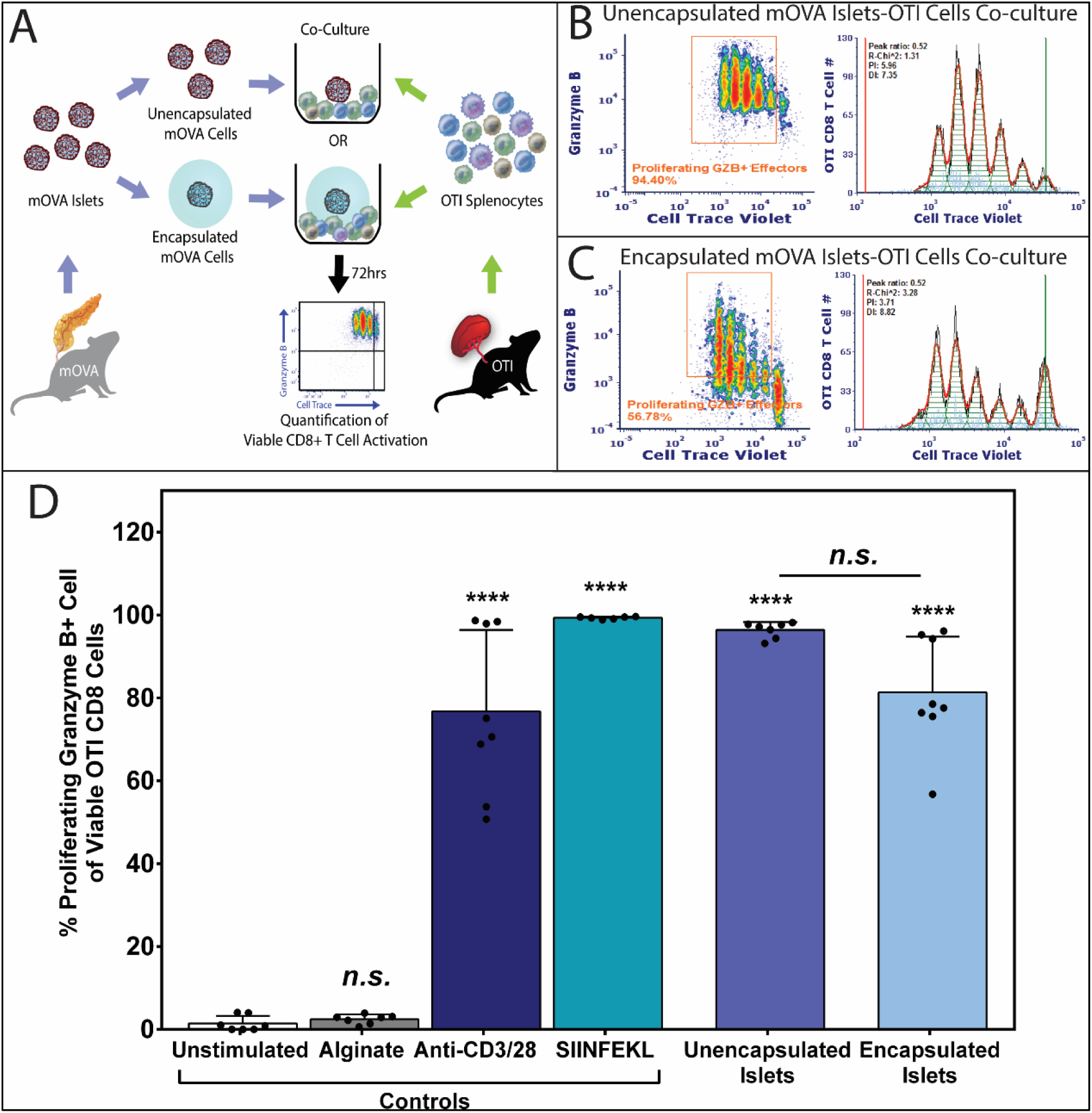
Alginate Encapsulated Islets Effectively Activate Antigen-Specific T cells via the Indirect Antigen Recognition Pathway. (A) Schematics of OTI splenocytes stimulated by unencapsulated or encapsulated mOVA islets for 72hrs. Antigen-specific CD8^+^ T cell activation was quantified using the method mentioned above (see **Figure S1**). Representative FCM data of OVA-specific proliferating granzyme B+ OTI CD8+ T cells effector and the respective proliferation fitting when 100,000 OTI responders stimulated by (B) 50 unencapsulated islets or (C) 50 complete encapsulated islets. Proliferation modeling was performed using FCS Express 6 software. Black line = histogram contour of raw data; orange line = fitted data; light blue points = noise events; green shaded area = area under the fitted curve; green marker = undivided marker and red marker = background marker. (D) Quantification of OVA-specific activated OTI CD8+ T cells, characterized as the percentage of proliferating and granzyme B+ CD8+ T effector cells. Data were shown as mean ± standard deviation of individual data points (N=3; n=9). Outliers were identified using Robust Fit with Cauchy estimate with multiplier K=2, using SAS JMP Pro v13.1.0. software. Tukey’s test was used for mean comparison. Statistical significance was determined as*\*\*\*\*p < 0.0001 or n.s. = not significant* with outliers removed for statistical analysis.

As the antigenicity of islets versus splenocytes may be disparate, the cell dosage and culture duration for islet experiments were evaluated. From these experiments, an antigen dose of 50 islets and an extended incubation of 72 hours were identified as optimal (**Figure S6**). Using these experimental conditions and the whole splenocyte repertoire, mOVA islets significantly activated OT1 CD8^+^ T cells. Similar to the splenocyte co-culture setting, polyclonal (anti-CD3/28 activator beads, 76.8 ± 19.7%) and monoclonal (SIINFEKL, 99.3 ± 0.3%) stimulators were used as positive controls, while unstimulated cell culture media (1.4 ± 1.8%) and the cell-free alginate hydrogel (2.4 ± 1.1%, Figure 6D) served as negative controls. Robust OT1 CD8^+^ T cell activation in response to both unencapsulated (96.4 ± 1.9%; Figure 6B,D) and encapsulated islets (81.3 ± 13.4%; Figure 6C,D) were observed. While the percentage of viable, granzyme B+ CD8^+^ T cells responding to unencapsulated or encapsulated mOVA islets was statistically equivalent (*p = 0.08*; post-hoc Tukey, Figure 6D), proliferation analysis demonstrated a delayed T cell response to the encapsulated islets, characterized by a lowered CD8^+^ T cell proliferation (PI = 6.06 ± 1.35) when compared to its unencapsulated counterpart (PI = 8.15 ± 1.72, *p = 0.04, t-test*; Figure 6B-C).

### 3.6 Indirectly activated T cells impair encapsulated pancreatic islets

With evidence of robust indirect adaptive immune T cell activation to alginate encapsulated cells, it was of interest to evaluate the impact of this immune cell activation on the underlying encapsulated cells. While the alginate barrier impedes direct interactions with the cytolytic CD8^+^ T cells, it is feasible that their activated state may contribute to a microenvironment unfavorable for islet viability and function. To address this hypothesis, islet microbeads were co-cultured with naïve OT1 splenocytes for 72 hrs and subsequently evaluated via live/dead viability imaging and glucose-stimulated-insulin-release (GSIR) functional assays.

When unencapsulated mOVA islets were co-cultured with OTI splenocytes, activated T cells aggressively attacked the islet, resulting in completely dissociated islets unamendable to subsequent assessments. For encapsulated mOVA islets, comparative assessments were made between islets co-incubated with OTI splenocytes or cultured alone for the same time period. As shown in Figure 7A&B, exposure of encapsulated islets to OTI splenocytes for 72 hrs resulted in detrimental impacts on peripheral islet cells, with a significant decline in viability when compared to untreated islets (*p<0.0001 respectively*; t-test; **Figure 8B**).

**Figure 7.**
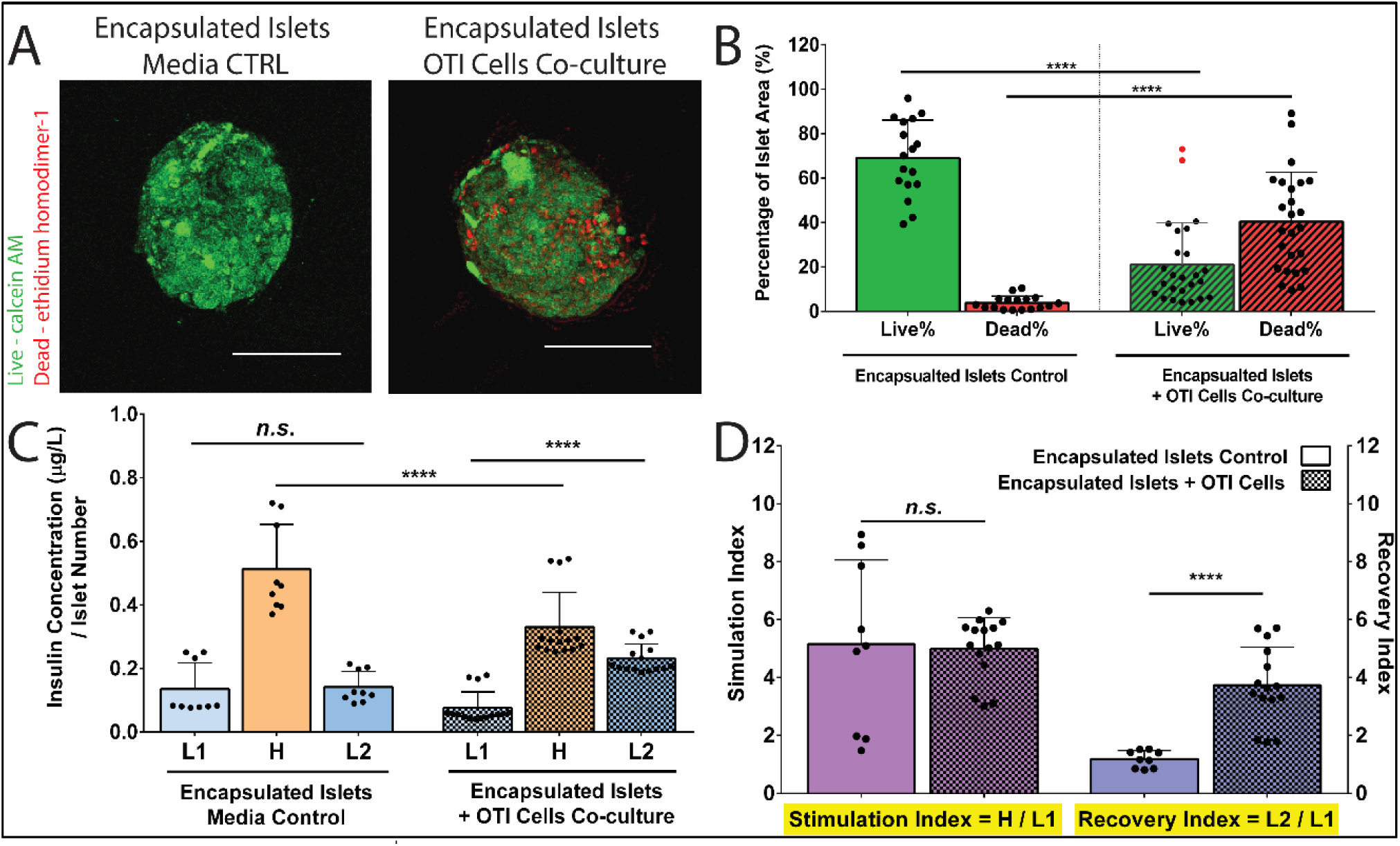
Indirectly Activated CD8^+^ T cells Impair the Viability and Function of Alginate Encapsulated Islets. (A) Representative Live/Dead images of alginate encapsulated mOVA islets following the 72hr co-culture with OTI splenocytes. Green= viable cells, Red = dead cells. Scale bars = 100 μm. Images analysis (B) quantified the % live cells and % dead cells of islets area (N=2; n=18-20). Outliers were identified using Robust Fit with Cauchy estimate with multiplier K=2, via SAS JMP Pro v13.1.0. software. Tukey’s test was used for mean comparison for viability quantification including the outliers. (C) Representative GSIR data of alginate encapsulated islets following 72hrs co-culture with OTI splenocytes or media control (n=9-15). Encapsulated islets were sequentially stimulated in 3 mM (Low1, L1); 16.7 mM (High, H), and another 3 mM (Low2, L2) glucose for 1hr respectively. Samples were collected after each hour stimulation, and the corresponding insulin level was measured by insulin Elisa. Insulin secretion was normalized by the number of the encapsulated islets for each sample. (D) Stimulation index (SI, the ratio of H/L1) and recovery index (RI, the ratio of L2/L1) of encapsulated islets after 72hr co-culture with OTI responder cells (n=9-15). Data were shown as mean ± standard deviation. Mean comparison was conducted using Tukey’s test. Statistical significance was determined as *\*\*p < 0.01; ****p < 0.0001 or n.s. = not significant*.

To evaluate functional impacts, insulin stimulation in response to a glucose challenge was assessed on the islet microbeads. In this test, islets were sequentially incubated in low (3 mM, L1), high (16.7 mM, H), and low (3 mM, L2) glucose for one hour respectively. This islet challenge permitted evaluation of insulin secretion in response to a high glucose challenge, followed by the termination of insulin release when glucose levels returned to basal conditions. A healthy islet response demonstrates a classic low-high-low insulin release profile, with a stimulation index (SI: H insulin release / L1 insulin release) > 1 and a recovery index (RI: L2 insulin release / L1 insulin release) ≈ 1 [51, 52]. For control islet microbeads, insulin release in response to glucose challenge was normal and responsive, with an SI = 5.15 ± 2.9 and an RI = 1.19 ± 0.3. When incubated with activated OTI CD8^+^ T cells, the function of encapsulated islets were negatively impacted, with decreased insulin production in response to high glucose stimulation (H_control_ = 0.51 ± 0.14 vs. H_T cell_ = 0.33 ± 0.10 μg/(L*IN); *p<0.0001*, t-test, Figure 7C). Impaired recovery of insulin release was also observed for islets microbeads challenged by extracapsular OTI CD8^+^ T cells, with an aberrantly high insulin release for the second low glucose incubation and a significant increase in RI (3.73 ± 1.32; *p<0.0001*, compared to control islet microbeads*, t-test*, Figure 7D).

To further delineate if the cytotoxicity observed in the encapsulated islet was attributed to soluble factors generated by activated T cells, culture media was procured from polyclonally activated OTI CD8^+^ T cells and added to encapsulated islet cultures. As summarized in **Figure S7**, viability and functional assessments illustrate substantial cell death and dysfunction imparted by the activated T cell media. Comparing the responses of unencapsulated and encapsulated islets, the alginate polymer dampens the negative impacts of activated T cell-conditioned media, with decreased cell death (**Figure S7A-E**) and improved GSIR (**Figure S7F-G**) compared to the unencapsulated islets; however, the negative impacts of the activated T cell-conditioned media on the underlying islets was profound, when compared to standard media controls (**Figure S7E&H**).

## Discussion

The concept of using polymeric encapsulating materials, particularly alginate, to prevent host recognition and rejection of foreign cells has been an area of cell-based therapy for decades. Despite numerous company ventures and clinical trials, the clinical translation of cellular encapsulation-based devices for the treatment of diseases, specifically for autoimmune diabetes, has resulted in disappointing outcomes, with poor functional efficacy and retrieved encapsulated implants containing significantly distressed cells [19, 25, 53–58]. There are multiple hypotheses as to what features contribute to the clinical failure of alginate encapsulated transplants, such as hypoxia and poor vascularization. Analysis of explants, however, suggests that host immune cells play a role [19, 25, 53–58]. Thus, elucidating the key features within encapsulated implants that instigate host immune responses is required to improve clinical outcomes.

As the encapsulating material serves as the interface between the implant and the host, most investigations into host immune responses have focused on the “biocompatibility” of the materials via characterization of host innate immune cells and their corresponding responses. It has been shown that common contaminants in alginate can activate macrophages via NF-κB pathway and promote DC maturation, as measured by the up-regulation of CD86 and HLA-DQ [59, 60]. Consequently, these activated innate cells direct classic foreign body responses that lead to fibrotic encapsulation [61, 62]. Modifications in alginate composition, purification, and final 3-D format have mitigated innate immune activation, resulting in decreased fibrotic responses to empty hydrogels and subsequently improving cell transplant outcomes in pre-clinical models [29, 63–65]. These mechanistic investigations, while important, focus primarily on one facet of the host immune response.

Animal transplant studies, however, indicate that host adaptive immune cells can sense, respond, and impact encapsulated cellular implants in an antigen-specific manner, supporting the theory that immune cells migrating to the implant site are likely not only reacting to the inert biomaterial [66–71]. For example, in rodent studies, the functional duration of encapsulated grafts is significantly reduced if the degree of antigen diversity between the donor and the recipient is elevated (i.e., allografts function longer than xenografts) or if T cells specific to the transplanted cell antigen are present (i.e. native mice function longer than mice primed/pre-exposed with antigen) [55, 67, 72]. T cell depletion and adoptive transfer studies implicate CD4^+^ T cells in the rejection of encapsulated xenograft transplants [67], with pharmaceutical interventions targeted at host T cell suppression, such as CTLA-Ig and anti-CD154 mAb, resulting in significantly improved encapsulated xenograft survival [31, 32, 73]. B cells must also sense the encapsulated graft, as *de novo* anti-xenograft antibodies emerge following the implantation of encapsulated xenogeneic cells [67, 68, 74]. Altogether, these transplant studies highlight the potentially deleterious impacts of the host adaptive response to encapsulated transplants.

The reliance on animal models to study immune activation pathways and screen new pharmaceutical targets, however, is not ideal. *In vivo* mechanistic studies characterizing adaptive immune responses to encapsulated cells are challenged by their inability to delineate host responses initiated by the non-specific factors (e.g., the implantation procedure, material compatibility, and the transplant site) from antigen-specific factors. Furthermore, characterization of the distinct roles of different immune cell components and antigen recognition pathways (e.g. direct versus indirect) towards microencapsulated cell cells, is difficult to ascertain *in vivo*. Finally, the substantial time and expense required for screening different encapsulation and pharmaceutical approaches in rodent models limit the further improvement of immunomodulatory biomaterial approaches. Developing a validated and effective *in vitro* platform that permits clear exploration of the interactions between host adaptive immune cells and encapsulated cell implants, particularly for identifying autoimmune-based therapies, could alleviate many of these challenges.

The *in vitro* platform developed herein was inspired by the classic mixed lymphocyte reaction (MLR), a simple *in vitro* assay that examines antigen-specific immune reactivity and histocompatibility. In the classic MLR approach, immune cells from two donor sources are co-cultured, where the T cell activation of the responder strain in response to the stimulator strain is quantified as the readout. While a powerful *in vitro* assay, a successful screening using MLR assay requires robust phylogenetic disparity of the tested strains and elevated frequency of reactive T cell precursors [75, 76]. Thus, when a classic MLR assay is used to explore naïve T cell reactivity to encapsulated cell grafts, results are typically weak and poorly reproducible due to insufficient reactivity and clonal specificity [53, 77–79]. For studying islets, this limitation may be further exacerbated by the poor proficiency of healthy, unstressed pancreatic beta cells at MHC-I or MHIC-II antigen presentation [75, 80, 81].

To overcome these challenges, an mOVA-OTI single-antigen model was employed. Stimulating mOVA cells provide the proficient surface presentation of OVA protein, as well as OVA peptide derivatives, such as SIINFEKL [82]. OTI T cells, which carry Vα2/Vβ5 receptors specific for OVA-derived antigen (SIINFEKL), render efficient T cell activation in response to encapsulated mOVA stimulators within a brief culture window (48 to 72 hours), permitting efficient screening [48]. In this study, alginate encapsulation completely inhibited the activation of purified OTI CD8^+^ T cells in response to mOVA cells, blocking direct donor-host contact and preventing direct T cell activation. Conversely, when the entire repertoire of OTI splenocytes was used as responder cells, robust T cell proliferation and effector function in response to the encapsulated mOVA cells was observed. These results indicate that CD8^+^ T cells can be activated by alginate encapsulated cells, when other immune cells, such as APCs, are present. These results are the first reported evidence, to our knowledge, that clearly demonstrate the impact of encapsulation on the activation of T cells via indirect recognition pathway *in vitro*.

To delve into the hypothesis that the indirect T cell activation by encapsulated cells is facilitated by host antigen presenting cells (APC), mechanistic studies using cross-presenting OTI CD8a^+^ DCs were conducted. CD8a^+^ DC was selected as the APC population of interest due to the MHC-I restricted features of the OTI model, which limited T cell activation solely to CD8^+^ T cells in lieu of the classic MHC-II APC and CD4^+^ T cell activation pathway [43, 83]. CD8a+ DCs also have an established role in apoptotic antigen uptake and presentation triggered by apoptotic and/or necrotic shedding antigens, a likely pathway for CD8^+^ T cell activation to encapsulated cell grafts [84, 85]. Our results found that the addition of CD8a^+^ DCs together with purified CD8^+^ T cell responder populations resulted in vigorous indirect CD8^+^ T cell activation in response to encapsulated mOVA cells. This activation was fully suppressed when the cross-presenting capacity of the DCs was inhibited via brefeldin A, further validating this host-donor contact-independent T cell activation pathway. Overall, this platform provides definitive evidence that CD8^+^ T cell activation in response to encapsulated cells persists via donor antigen cross-presentation by CD8a^+^ DCs. Of note, other immune cells, e.g., CD4^+^ T cells, macrophages, and B cells, may also play a role in the collective adaptive immune responses to encapsulated grafts and are the subject of future studies.

This encapsulation assay can subsequently be used to intimately study rejection pathways and optimize new biomaterials for encapsulated cells, as well as to screen complementary immunomodulatory therapies. As an example of this utility, the role of antigen dosage on antigen-specific T cell activation was characterized. OTI CD8^+^ T cell activation in response to mOVA cell-derived antigens was generally depressed by the encapsulating hydrogels, both in terms of proliferation index and generation of effector T cells. Elevation of antigen dosage via increased total cell number, however, resulted in increased activation. Of interest, increasing encapsulation cell density, while maintaining a constant total number of mOVA cells, resulted in increased T cell activation. These results indicate that the number of cells and the manner by which they are loaded into an encapsulation system may impact downstream adaptive immune responses. These correlations also implicate additional activator factors distinct from the total cell number. Increasing the cell loading density within the microbead results in an increased percentage of stressed and/or nutritionally deficient cells. Distressed cells are known to release of instigative agents, such as damage-associated molecular patterns (known as DAMPS, e.g., high mobility group box-1) and inflammatory cytokines (e.g., IL-8, TNF-α), which can easily permeate out of the hydrogel [69, 70]. Co-delivery of these agents with antigen may exacerbate adaptive immune cell activation. As such, efforts that seek to improve nutrient availability to the grafts, via improved oxygenation, vascularization, and/or transplant sites, can be leveraged to improve the efficacy of the encapsulated cell grafts[86, 87].

To apply this *in vitro* platform to a disease model, encapsulated pancreatic mOVA islets were investigated. Using mOVA islets as the stimulatory graft, significant CD8^+^ T cell activation was observed via the indirect antigen presentation pathway. Despite the impairment of direct cell contact between the T effector cells and islet grafts, indirectly activated T cells imparted detrimental effects to the encapsulated cells, with cell death observed at the periphery of the islet. This cell death pattern implicates an external cytotoxic stimulus, as opposed to insufficient nutritional availability. Islet function was also impacted, with decreased insulin release under a high glucose challenge and a reduced capacity to shut down insulin release following return to non-stimulatory glucose levels. While the mechanism of persisting high insulin release following exposure to stimulatory glucose is not fully understood, it has been attributed to elevated oxidative stress or ionic channel leakage, which can be induced by exposure to inflammatory cytokines [88–91]. Supplemental experiments using only activated T cell-conditioned media resulted in similar impairment to the encapsulated islets, further confirming that T cell-derived diffusible cytotoxic solutes, such as TNF-α (17.5kDa) or IFN-γ (16.9kDa), likely contributed to the observed cytotoxicity [92]. Islet cytotoxicity was mitigated by the presence of the encapsulating hydrogel, as evidenced by elevated peripheral cell death for unencapsulated islets exposed to the sample conditioned media. While the protective nature of alginate from peripheral cytotoxic agents has been postulated, this study highlights its protective, albeit incomplete, effects [70]. Further elucidation of the specific diffusing cytokines responsible for this cytotoxicity will inform improved encapsulation approaches. Overall, the successful translation of the mOVA-OTI model to the study of encapsulated islet demonstrated the flexibility and versatility of our antigen-specific *in vitro* platform, which allows the screening of stimulator cells of different tissue types for different disease models and cell therapies.

While this assay provides a powerful *in vitro* platform for immune screening, it is fully recognized that the efficiency of this assay is skewed by the high affinity of the transgenic Vα2/Vβ5 T cell receptors to OVA peptides, as well as the elevated antigen presence associated with the mOVA cells [93]. *In vivo*, the timeline of T cell activation would be extended, given the need for antigen uptake and processing by APCs, trafficking, antigen matching, and clonal expansion. However, the ease by which the APC-mediated T cell activation occurs in our clonally specific platform permits clear evidence that the indirect antigen pathway is fully functional for encapsulated grafts. In addition, this clonal specificity serves to mimic the clinical situation where indirect antigen presentation and T cell activation are dominant, such as in established autoimmunity [35, 75, 94].

With the establishment of this antigen-specific *in vitro* platform for the detailed characterization of host immune responses to encapsulated cell grafts, future work will focus on exploring the roles of non-cross-presenting APCs and CD4^+^ T cells in indirect T cell activation, *e.g.*, by adjusting the responder cells population or using MHC-II restricted OVA-specific responders [95]. Moreover, the presented antigen-specific *in vitro* platform will be leveraged to screen immunomodulatory biomaterials targeted to dampen the unique adaptive immune cell pathways activated by encapsulated cells.

## Conclusion

In this study, we successfully developed an efficient, antigen-specific *in vitro* co-culture platform with robust reproducibility to study host adaptive immune responses to hydrogel encapsulated cell grafts. Leveraging our platform, we conclude, with detailed experimental evidence at the cellular level, that alginate microencapsulation effectively blocks the host-to-donor, contact-dependent, direct T cell activation, while the host APCs-mediated indirect T cell activation to the encapsulated cell grafts is preserved. This resulting indirect T cell activation can impart detrimental effects on the encapsulated cell grafts via the diffusion of soluble cytotoxic agents. Overall, this reported mOVA-OTI *in vitro* platform provides ease of immune screening for biomaterial-based encapsulation approaches. The findings with this platform can inspire the design of bioactive encapsulating materials targeting the suppression of adaptive immune recognition for improved clinical outcomes for cell replacement therapy.

## Acknowledgments

This work was supported by NIH grant DK100654. We thank Drs. Craig Moneypenny and Andria Doty of the University of Florida ICBR Cytometry Core and Howie Seay of UF Center for Immunology and Transplantation for cell sort assistance; Chad Rancourt of Animal Care Services of the UF for the assistance of animal breeding; and Cecilia Cabello of the University of Miami for the assistance of animal genotyping. We also thank all members of the Stabler lab for their collective assistance in animal care and monitoring, as well as islet isolations.

## Author contributions

Y.L., A.W.F., E.Y.Y., A.L.B. and C.L.S. developed the concept. Y.L., A.W.F., E.Y.Y, and C.L.S. designed the study and analyzed data. Y.L., A.W.F., C.S., Y.R., E.Y.Y, and M.M.S. conducted experiments, data collection and analysis. I.M.L. and M.M.S. assisted in the conduction and optimization of islet isolation. All authors helped in writing the manuscript.

## Abbreviation

APC: Antigen Presenting Cell
DC: Dendritic cell
DI: Division index
DAMPS: Damage-associated molecular patterns
ELISA: Enzyme-linked immunosorbent assay
FCM: Flow cytometry
GFP: Green Fluorescent Protein
GSIR: Glucose stimulated insulin response
HBSS: Hank’s Balanced Salt solution
mAb: Monoclonal antibody
MHC: major histocompatibility complex
MLR: Mixed lymphocyte reaction
mOVA: membrane-bound ovalbumin
OVA: Ovalbumin
PBS: Phosphate buffered saline
PI: Proliferation index
RI: Recovery index
SI: Stimulation index
T1D: Type 1 Diabetes

