## Supporting Documents for "*In Vitro* Platform Establishes Antigen-Specific CD8^+^ T Cell Cytotoxicity to Encapsulated Cells via Indirect Antigen Recognition"

**SUPPORTING TABLES**

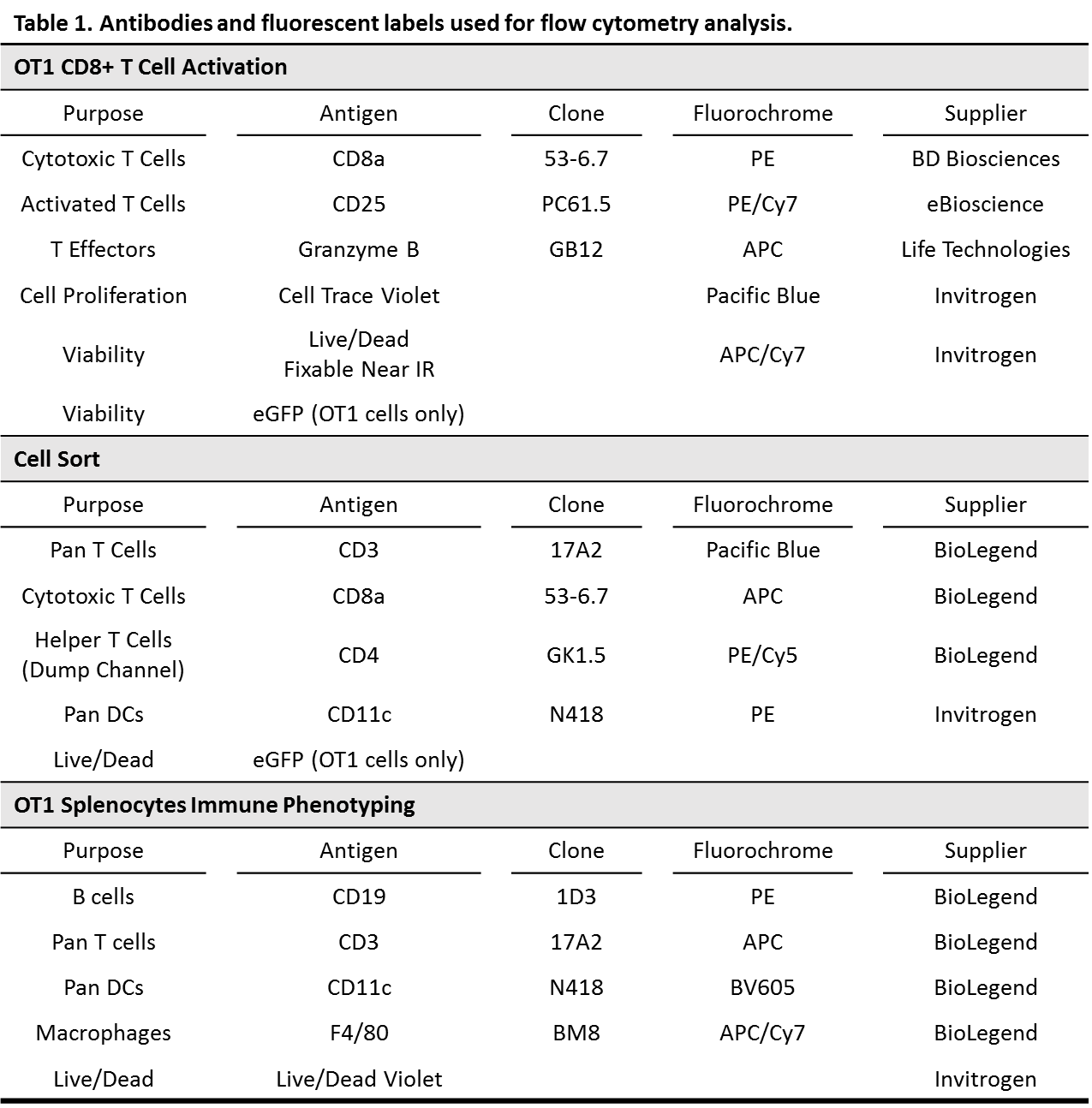

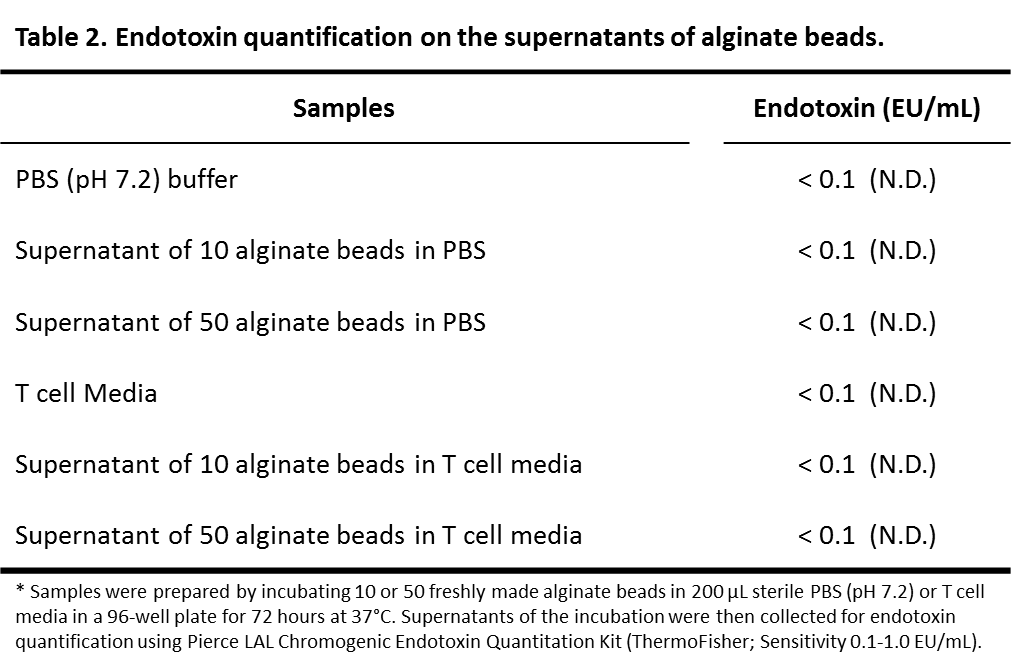

**SUPPORTING FIGURES**

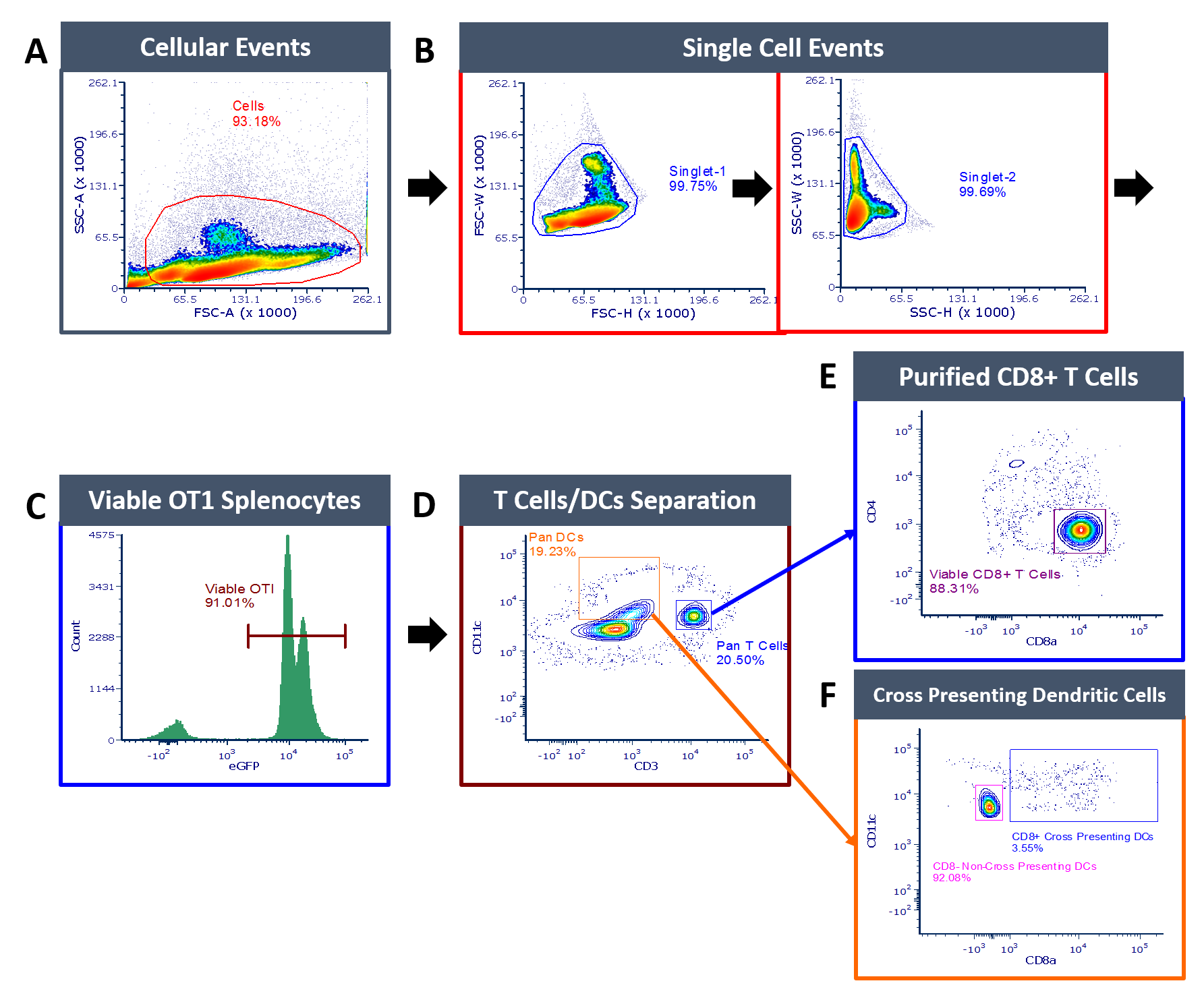

**Figure S1. Representative Cell Sorting Gating Strategy used to Purify OTI CD8^+^ T cells and OTI CD8^+^ Cross-presenting Dendritic Cells.** Freshly isolated OT1 splenocytes were stained and used for cell sorting. Cell debris and aggregates were excluded by serially gating as depicted in (A) and (B). Then viable OTI splenocytes were gated out base on (C) eGFP expression. Purified OTI CD8^+^ T cells were identified and collected as the CD3+CD11c-CD8+CD4- population, as shown in (D) and (E). The cross-presenting dendritic cells from the same splenocyte isolation were isolated as the population CD3-CD11c+CD8+ shown in (D) and (F). Gating boundaries were determined based on fluorescence-minus-one (FMO) controls and the unstimulated control with proper compensation. Three cell sorts were independently performed using BD FACSAria II/III Cell Sorter with efficiency over 98%.

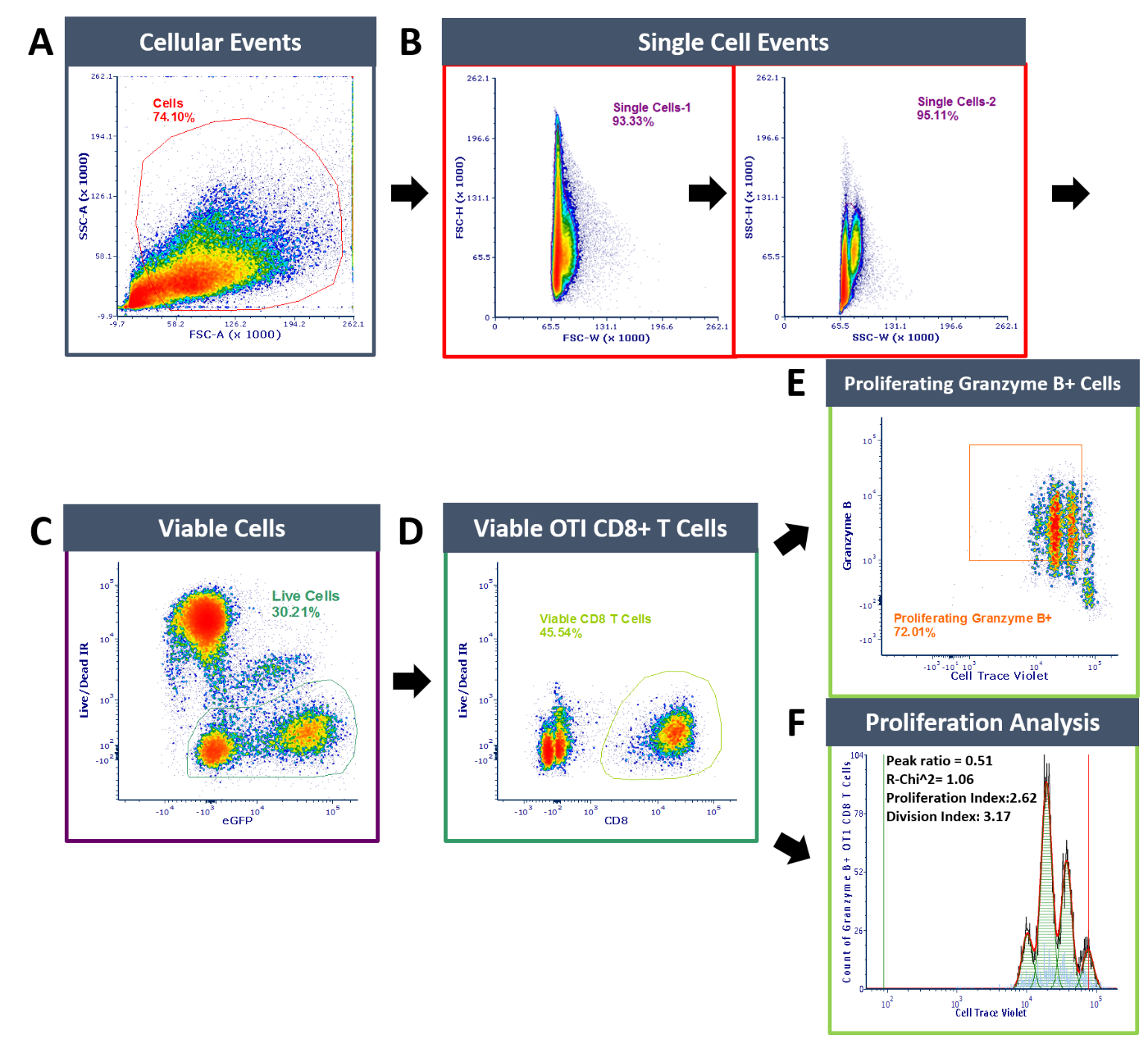

**Figure S2. Representative flow cytometry gating strategy for enumeration of granzyme B^+^ OTI CD8 effectors.** Sequential gating strategies were applied to quantify OTI CD8^+^ T cell activation as followed. (A) Set a polygon gate to identify the lymphocyte population and exclude debris. (B) Exclusion of cell doublets and aggregation. (C) Live/Dead IR negative cells were identified as viable cells. (D) Viable CD8^+^ T cells were identified. (E) Proliferating (excluding generation 0 of cell trace violet signal) granzyme B expression on viable OTI CD8^+^ T cells; (F) Proliferation analysis on viable OTI CD8^+^ T Cells, where black line = histogram contour of raw data; orange line = fitted data; light blue = noise events; green shaded area = area under fitted curve; green marker = undivided marker and red marker = background marker. Proliferation analysis was performed using FCS Express 6.05 software. The boundary of the gates is determined by isotype controls, fluorescence-minus-one (FMO) controls and the unstimulated control.

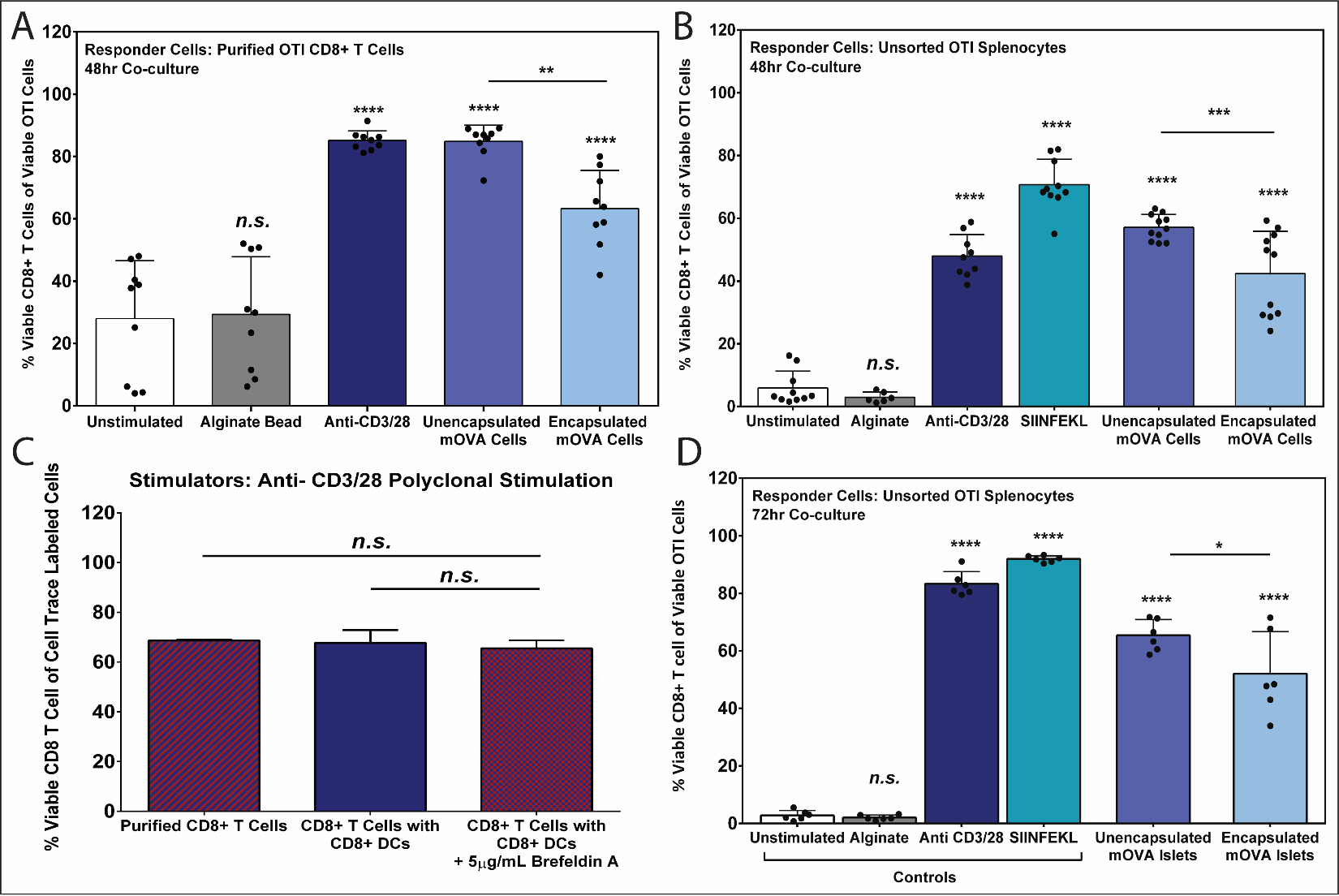

**Figure S3. Responder OTI CD8^+^ T cell viability after *in vitro* co-culture assay.** Responder cell viability (%) was quantified via flow cytometry. A-B) Responder OTI CD8^+^ T cell viability, expressed as %, from (A) purified OTI CD8^+^ T cell (N=3; n=9) and (B) unsorted OT1 splenocytes co-cultured (N=4; n = 6-11 each group) following co-culture with the designated stimulant (x-axis) for 48hrs. (C) CD8^+^ T cell viability (%) of the responder pool, designed by x-axis, following stimulation by anti-CD3/28 activator beads for 48 h (n = 3). (D) CD8^+^ T cell viability (%) for unsorted OTI splenocytes co-cultured with designated stimuli for 72 h (N=2; n=6). Data were shown as mean ± standard deviation. One-way ANOVA with Tukey post-hoc. **p < 0.05; **p < 0.01; ***p < 0.001; ****p < 0.0001 or n.s. = not significant*.

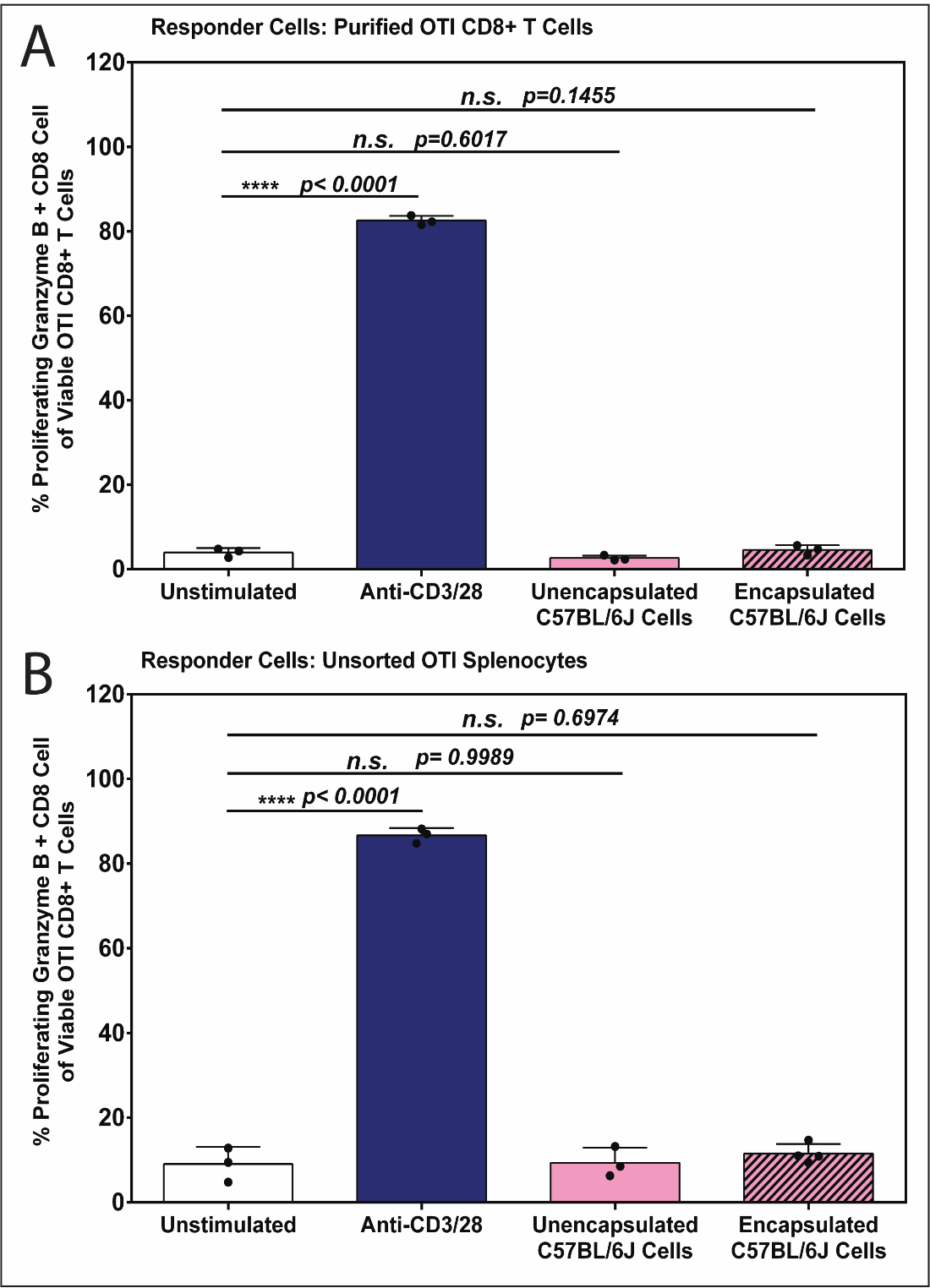

**Figure S4. CD8^+^ T Cell Activation of Purified OTI CD8^+^ T Cell and Unsorted OTI Splenocyte is Antigen-Specific by mOVA cells.** Splenocytes isolated from C57BL/6J mice (MHC-matched; H2K^b^), in either unencapsulated or alginate encapsulated form, were used as unspecific stimulator cells to co-culture with OTI splenocytes, to validate antigen specificity of the system. OTI CD8+ T cells activation was quantified by the percentage of viable proliferating granzyme B positive CD8^+^ effectors via flow cytometry analysis post 48hr stimulation. Data were shown as mean ± standard deviation. Tukey’s test was used for mean comparison (n=3). Statistical significance was determined as *****p < 0.0001*; *n.s.* = not significant; when compared to the unstimulated control group (T Cell Media).

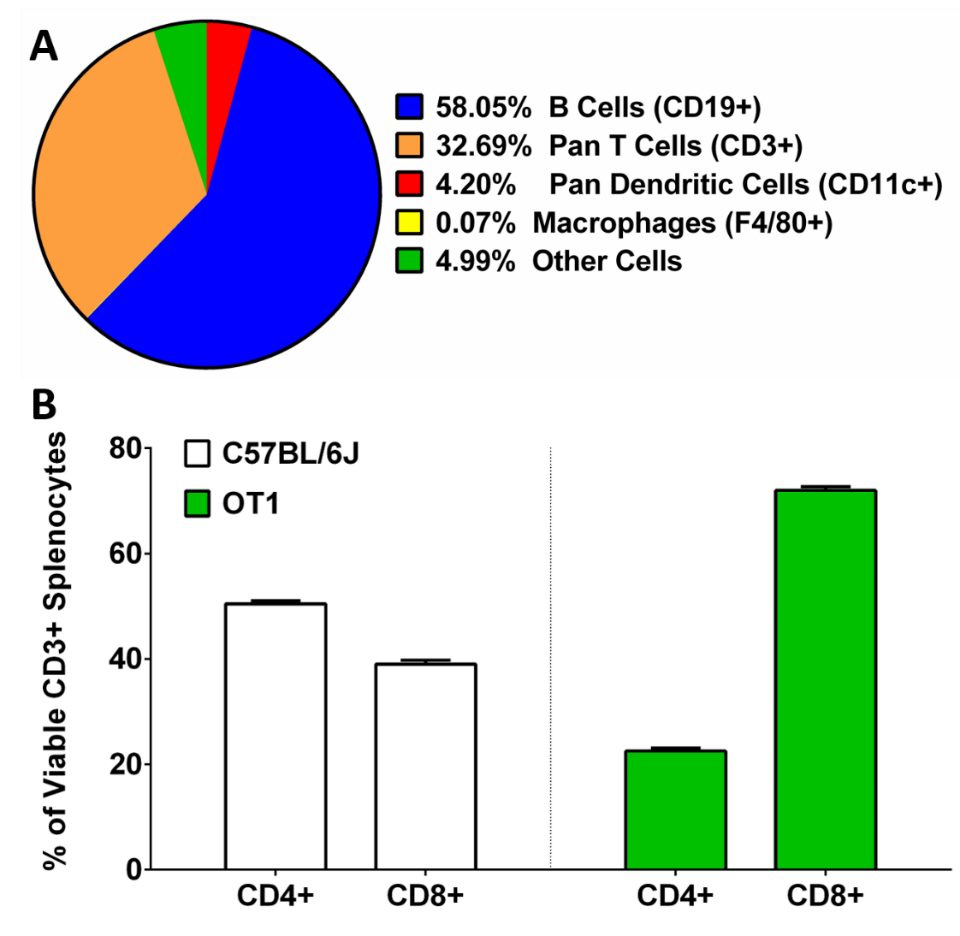

**Figure S5. Cellular Component of OTI Splenocytes.**  (A) Immune phenotyping of fresh isolated OTI splenocytes via flow cytometry. Viable cells were identified using Live/Dead violet staining (Invitrogen). The frequency of different cell populations was determined by the immune labeling of anti-mCD19-PE for B cells, anti-mCD3-APC for pan T cells, anti-mCD11c-BV605 for pan DCs and anti-mF4/80-APC/Cy7 for macrophages (n=3) (see **Table1**). (B) CD4^+^ and CD8^+^ T lymphocyte distribution of unsorted splenocytes isolated from C57BL/6J and OTI mice. The frequency of CD4^+^ and CD8^+^ cells were quantified within the viable (Live/Dead staining-) T cell (CD3+) population. Data were shown as mean ± standard deviation (n=3).

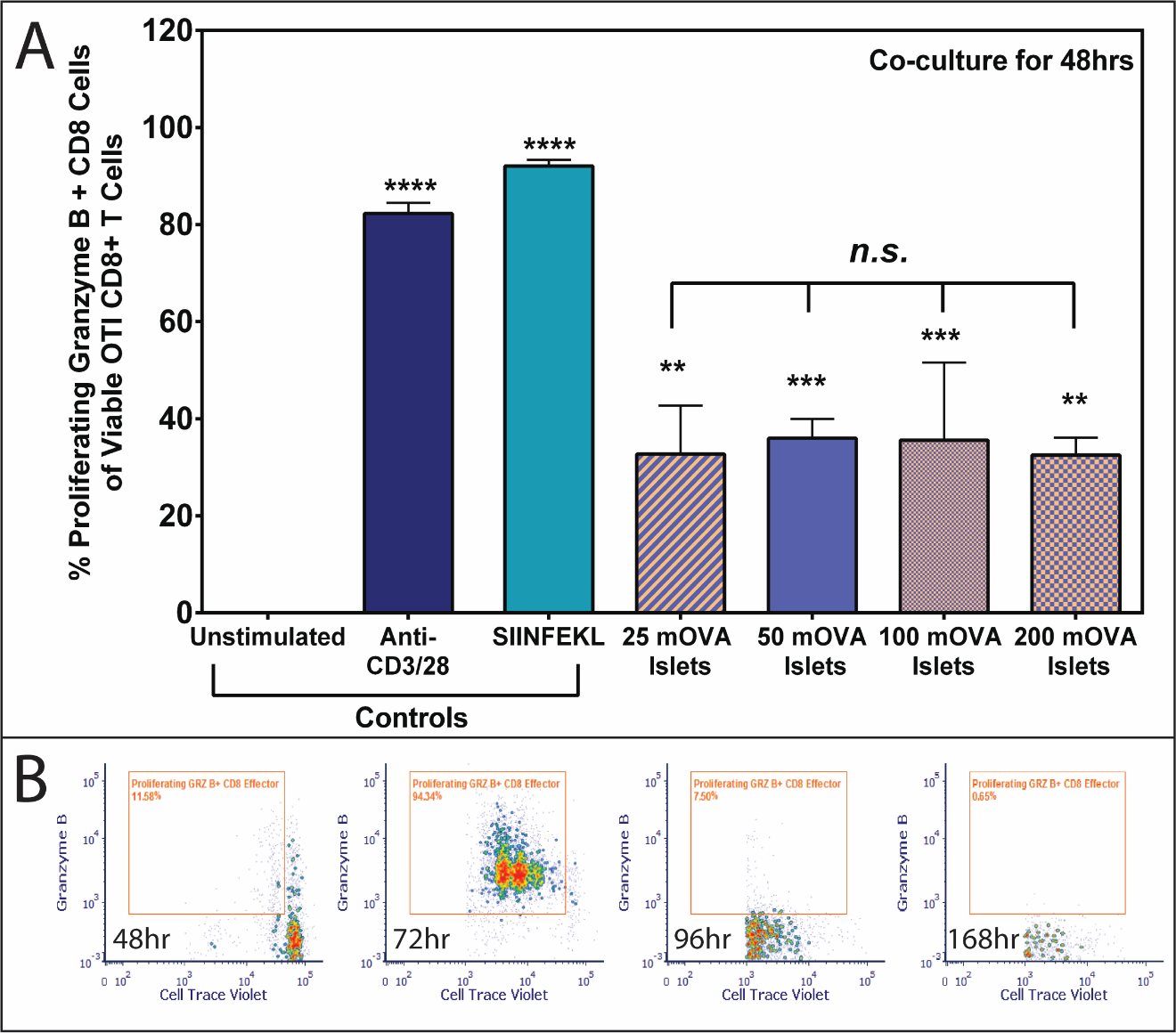

**Figure S6. Antigen Titration and Time Course of OVA-specific OTI CD8^+^ T cell Activation by mOVA islets.**  (A) Summary of OTI CD8+ T cells activation by unencapsulated mOVA islets titration for 48 hours co-culture (n=3). Control groups were unstimulated controls (complete medial), 0.1μM SIINFEKL peptide and anti-CD3/28 activator beads. Data were shown as mean ± standard deviation. Mean comparison was conducted using Tukey’s test. Statistical significance was determined as ***p < 0.01; ***p < 0.001; ****p < 0.0001 or n.s. = not significant*. (B) Representative time course FAC data of the OVA-specific OTI CD8^+^ T cell activation (see **Figure S1** for gating) stimulated by 50 purified mOVA islets (N=2, n=3-6).

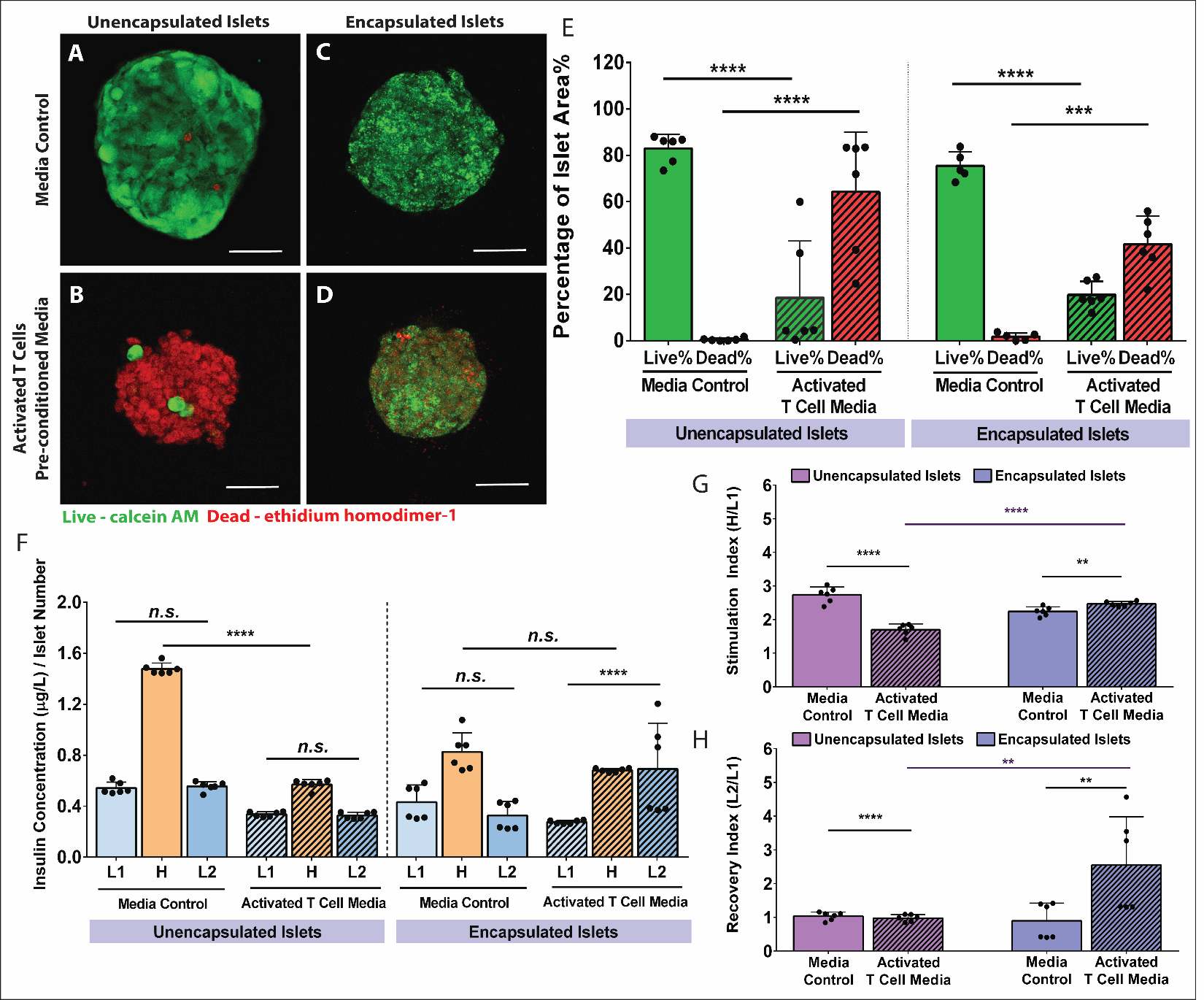

**Figure S7. Encapsulated C57BL/6 Islets Cytotoxicity by Activated T Cell-Derived Cytokines.** Representative Live/Dead images of unencapsulated (A-B) and alginate encapsulated (C-D) C57BL/6 islets following incubation with CMRL media (A, C) or activated OTI T cell media (B, D). Green= viable cells, Red = dead cells. Scale bars = 100 μm. Images analysis (E) quantified the percentages of live cells and dead cells of islets (n=6). (F) Representative GSIR data of unencapsulated and alginate encapsulated islets following a 48hr challenge by activated T cell-derived pre-conditioned media. L1 and L2 = 3mM glucose; H = 16.7 mM glucose (n=6). (G) Stimulation index (SI, the ratio of H/L1) and (H) recovery index (RI, the ratio of L2/L1) of islets challenged by controls and activated T cell pre-conditioned media. Data were shown as mean ± standard deviation. Mean comparison was conducted using Tukey’s test. Statistical significance was determined as **p < 0.05; **p < 0.01*; ****p < 0.001; ****p < 0.0001* or *n.s. = not significant*.

**Supporting Methods**

- **Endotoxin Quantification**

Endotoxin levels of the alginate were tested by incubating 10 or 50 freshly made cell-free alginate beads in 200 μL sterile PBS (pH 7.2) or T cell media in a 96-well round-bottom plate (Corning) for 72 hours at 37°C. The supernatant of the incubation was then collected by centrifugation at 300 ×g for 10 min. PBS buffer and T cell media were used as blank controls. The levels of endotoxin were quantified using Chromogenic LAL kit (Thermofisher; Sensitivity 0.1-1.0 EU/mL) following the manufacturer’s instructions. Three independent experiments were conducted.
